# Phenotypic Approaches to T Cell Activation: A Comparative Mathematical Modeling Study

**DOI:** 10.1101/2025.04.07.647688

**Authors:** Yogesh Bali, Alan D. Rendall

**Affiliations:** Institut für Mathematik, Johannes Gutenberg-Universität, Staudingerweg 9, 55099 Mainz

**Keywords:** T cell activation, kinetic proofreading, phenotypic models, mathematical analysis, sensitivity analysis

## Abstract

T cells use their T cell antigen receptors (TCRs) to recognize peptides presented by major histocompatibility complex molecules (pMHC). These peptides may be low-affinity self-peptides or high-affinity foreign peptides from pathogens. Despite recognizing a broad range of affinities, TCRs trigger significant immune responses only to strongly binding foreign peptides. The mechanisms enabling TCRs to distinguish diverse antigens with high sensitivity remain a key focus of research.

Our goal is to analyze mathematical models of T cell activation for their ability to replicate key experimental features like optimal response, specificity, sensitivity, and antigen discrimination. We analyzed nine models using mathematical and numerical methods to examine their solutions, responses, and parameter sensitivity.

We found that in all models, except kinetic proofreading with negative signaling, solutions converged to a unique steady state. Most response functions defined by ligand concentration and dissociation time showed an optimum value, except for the Occupancy, KPR, and stabilizing activation chain models. Models like KPR with negative feedback, limited/sustained signaling, and incoherent feedforward loops effectively replicated the key features of specificity, sensitivity, and antigen discrimination. Our sensitivity analysis identified phosphorylation rate as a key parameter influencing most model outcomes. This study highlights the strengths and limitations of current T-cell activation models, suggesting improvements to enhance their predictive accuracy in future research.

## 1 Introduction

T cells continuously navigate through lymph nodes, spleen, and various lymphoid tissues while actively searching for antigens. During this exploration, T cells survey the surfaces of specialized cells, particularly antigen-presenting cells (APCs), aiming to identify and react to foreign or irregular molecules, known as antigens, presented on these cell surfaces. Through their T cell receptors (TCRs), T lymphocytes interact with antigens presented by MHC molecules on the surfaces of APCs in the form of peptide-MHC complexes (pMHC). This ongoing scanning process plays a pivotal role in the immune system’s ability to detect and counter infections or aberrant cells, including those linked to conditions such as cancer [1].

Understanding the process of T cell activation is essential for understanding how immune responses are initiated at the cellular level. This interaction serves as a cornerstone of the adaptive immune system, initiating intricate signaling pathways within cells. These pathways trigger the activation of specific genes crucial for various T cell mediated responses, including proliferation, cytokine release, and cytotoxic actions. An in-depth analysis of these phenomena sheds light on the mechanisms by which the immune system is activated and functions to combat infections and diseases [2].

Although the T cell pool includes a diverse collection of unique TCRs capable of recognizing a broad spectrum of antigenic peptides, only a small fraction of an individual’s projected total T cell population (ranging from 10^11^ to 10^12^) [3, 4] could effectively recognize a specific agonist pMHC. Given that the TCR is the only structure on the T cell responsible for detecting antigens, it is important to understand how the TCR discriminates between self and foreign ligands. To explore TCR-pMHC interactions numerous researchers have conducted extensive experiments. These investigations have led to the development of several theories that tried to explain the mechanisms underlying activation of T cells.

One of the first mathematical frameworks proposed to explain T cell activation is the occupancy model. This model posits that activation of T cells is directly related to the quantity of TCR-pMHC complexes formed during the binding event. Based on this premise, even low-affinity pMHC molecules should theoretically trigger T cell activation if present in sufficiently high concentrations. However, experimental findings contradict this assumption, demonstrating that an increased concentration of low-affinity pMHC fails to activate T cells effectively [5]. In contrast, pMHC complexes with relatively higher affinity are capable of activating T cells even at low concentrations [5, 6].

To explain this affinity based discrimination, the kinetic proofreading (KPR) model was proposed [7]. This model suggests that only pMHC complexes that stay attached to TCRs for a sufficient duration can achieve a signaling competent state. During this time the TCR-pMHC complex undergoes mechanical processes such as the initial dimerization of receptors, the phosphorylation of multiple tyrosine residues on the TCR complex, and the subsequent recruitment and activation of ZAP-70. In fact, experimental findings have revealed differences in the early phosphorylation patterns of molecules like p21/p23 and in the recruitment and activity of ZAP-70 in response to pMHCs of varying potencies [8, 9]. Therefore, low-affinity pMHC cannot activate T cells because the total dissociation time of low-affinity pMHC complexes is shorter compared to high-affinity pMHC. Thus, the KPR model successfully explains the specificity of TCR-pMHC interactions.

Since the KPR model considered the threshold time as the key factor for TCR-pMHC interactions to reach the signaling state, it implies that an increase in dissociation time enhances specificity. Experimentally, this effect is evident when comparing pMHC ligands with dissociation times of 5 seconds and 1 second. A ligand with a 5-second dissociation time is 55 times more likely to stay bound beyond the threshold, thereby successfully initiating a signal [6]. However, this gain in specificity comes at the cost of sensitivity. An extended threshold time means that the fraction of pMHC bound decreases exponentially, which in turn diminishes the sensitivity of the TCR-pMHC interaction. This trade-off highlights the delicate balance between achieving high specificity and maintaining sensitivity in TCR-pMHC interactions.

Moreover, since antigenic pMHCs on the APC surface are scarce, the number of TCRs that can bind simultaneously is limited [10]. Studies with T cells having two different receptors show that when one TCR is activated, it does not affect nearby TCRs. This suggests that each TCR must directly interact with a pMHC to be activated. Further experiments confirmed that TCR activation happens independently for each receptor [11]. In accordance with this, experiments have shown that even a small number of pMHC complexes can activate many TCRs by binding to them one after another [12], with individual pMHCs sequentially activating numerous TCRs. These findings, along with the observations that TCR-pMHC interactions are low-affinity and rapidly dissociating, support the idea that each pMHC ligand can quickly attach and detach from a TCR, enabling continuous TCR engagement and sustained signaling. The serial triggering model suggests that swift pMHC release from previously engaged TCRs is necessary for sequential TCR engagement, potentially conflicting with the long bond lifetimes posited by the kinetic proofreading model. To reconcile this, an optimal dwell time concept was proposed [13], indicating that intermediate half-lives are essential for effective downstream signaling. The experimental backing for this model comes largely from studies employing mutated TCR panels [14]

To study TCR-pMHC interaction properties discussed above, researchers conduct dose-response experiments by exposing T cells to controlled pMHC doses [6]. These experiments help identify key factors influencing T cell activation, including TCR-pMHC affinity [15, 16], dissociation time [13, 17], kinetic segregation [18, 19], conformational changes [20], and the catch bond phenomenon [21, 22].

Furthermore, certain experiments highlight a correlation between dissociation time and potency (*EC*_50_, the pMHC dose needed for half the maximum response) as well as *E*_max_ (the peak response to a pMHC) [6, 23]. These factors significantly impact T cell activation, prompting various researchers to develop models that elucidate the extent and impact of these parameters. In addition, there is an on-going effort to create models that comprehensively explain the main phenotypic characteristics of T cell activation.

Our study aims to investigate the key characteristics of TCR-pMHC binding with three major objectives: (1) to mathematically analyze the behavior of solutions within selected models, (2) to identify models capable of reproducing critical features—such as antigen discrimination, specificity, sensitivity, and optimal response—based on biologically and experimentally established parameter values, and (3) to determine the parameters that most significantly impact model outcomes. To achieve these objectives, we analyzed various phenotypic models of TCR activation, focusing on those that can replicate these characteristics under specific parameter conditions. Our analysis includes nine models: seven are grounded in established experimental findings and analyses, while two are newly constructed by combining elements of existing models to explore additional outcomes.

We began by mathematically analyzing the existence and uniqueness of steady states in various models using the zero deficiency theorem [24]. This approach allowed us to rigorously confirm that all models, except the KPR model with negative signaling, possess unique and stable steady-state solutions. On the other hand the KPR model with negative signaling exhibits multiple steady states, as demonstrated by Rendall and Sontag [25]. These findings provide a robust foundation for further analysis and application [26, 27]. We mathematically calculated the response or activation of the models and plotted response as a function of ligand concentration and dissociation time, using parameters from four experimental sources [28, 29, 30, 31]. To ensure a comprehensive assessment, we tested the models with the mean, minimum, and maximum parameter values. Our key findings highlight the response optimum in the kinetic proofreading (KPR) model under both limited and sustained signaling conditions. In the negative signaling model, we observed an optimal response with respect to dissociation time; although not highly pronounced, this aligns with predictions made earlier [25]. Models incorporating incoherent feedforward (IFF) motifs exhibited an optimal response to both ligand concentration and dissociation time; however, at lower ligand concentrations, the response did not reach an optimum with respect to dissociation time. Notably, the KPR model with IFF and limited signaling exhibited a unique bimodal response as a function of dissociation time. Furthermore, the KPR model with both limited and sustained signaling demonstrated an optimum response with respect to both dissociation time and ligand concentration.

Building on findings from [6], we calculated the response function for all models, focusing on maximum response (*E*_max_) and half-maximal effective concentration (*EC*_50_) as key indicators of T cell activation mechanisms. We plotted *E*_max_ and *EC*_50_ curves in relation to dissociation time. The *E*_max_ plots revealed optimality for the KPR models with limited signaling, KPR with limited and sustained signaling as well as for the IFF models, while *EC*_50_ consistently showed a correlation with dissociation time across all models.

Given the high sensitivity of our results to parameter selection, we performed a stability analysis using Latin Hypercube Sampling-Partial Rank Correlation Coefficient (LHS-PRCC) [32] to identify the parameters with the greatest impact on model behavior. Our analysis revealed that the phosphorylation off-rate and phosphatase efficiency are the key drivers of the response function across most models. Additionally, we examined the experimental methods used to derive parameters for phenotypic models, comparing 2D and 3D approaches to highlight differences in their processes and outcomes. In the discussion, we explore potential strategies to enhance model robustness and address the limitations of existing models. This study provides a comprehensive assessment of model stability and dynamics, particularly in relation to ligand concentration and dissociation time across different configurations.

The remainder of this paper is organized as follows: Section 2 introduces the models, discusses the background of parameter values, and provides qualitative predictions alongside the plots of our numerical simulations. Section 3 delves into the motivations, characteristics, and comparative analysis of each model in relation to previous research. In that section we have also validated the assertions made for the case of KPR with negative signalling in [25] regarding the non-monotonic nature of the response as a function of dissociation time. Section 4 presents the LHS-PRCC analysis, identifying key parameters that most significantly influence model responses. Section 5 offers a comparative analysis of 2D and 3D experimental techniques used for parameter estimation, examining their impact on model accuracy. Finally, Section 6 provides a discussion of the results and outlines potential directions for future research.

In the appendix section, Appendix A presents the plot of the response function versus dissociation time and ligand concentration for the minimum parameter values. Appendix B contains the proof for the analysis of the late-time behavior of the models: Sections B.1 to B.7 provide the proof for the solutions of the models converging to a unique steady state, while Section B.8 discusses the non-monotonic response function observed in some models and its analysis. Appendix C provides detailed calculations for determining the response function, *E*_max_, and *EC*_50_ for all of the models.

## 2 Model Foundations: Definitions and Key Properties

Our analysis of phenotypic models of TCR activation focused on identifying which models can replicate key experimental characteristics such as specificity, sensitivity, antigen discrimination, and dose-response optima under specified parameters. We examined nine models, seven grounded in established experimental data and two developed by combining existing models to explore alternative outcomes. The models we considered are

1. Occupancy Model
2. Kinetic Proofreading
3. Kinetic Proofreading with Limited Signaling
4. Kinetic Proofreading with Sustained Signaling
5. Kinetic Proofreading with Negative Feedback
6. Kinetic Proofreading with Stabilizing Activation Chain
7. Kinetic Proofreading with Incoherent Feed Forward Loop
8. Kinetic Proofreading with Limited Signaling and Incoherent Feed Forward Loop
9. Kinetic Proofreading with Limited and Sustained Signaling

Of these nine models, the first four (1), (2), (3), and (4) were reviewed by [28]; the KPR model with negative feedback (5) is described in [29]; and the KPR with activation chain stabilization (6) comes from [30]. Model (9) combines elements from (3) and (4), while the KPR with limited signaling and incoherent feedforward loop (8) is based on [31]. Finally, (7) is constructed by linking (2) with an incoherent feedforward loop.

There are a number of relations between these models. The notations used in describing these are defined in later sections. Setting *ξ* = 0 in Model 3 gives Model 2. In the modified version of Model 3 the equation for *C*_*N*+1_ decouples and this variable tends to zero for *t* → ∞. Thus if the dynamical properties of solutions of Model 3 are known those of solutions of Model 2 can be concluded. Setting Ω = 0 in Model 4 means that adding the last two equations shows that the *C*_*i*_ define a solution of Model 3. Similarly, setting *ξ* and Ω to zero in Model 9 produces a solution of Model 2. Model 2 is a special case of Model 6 obtained by setting some of the parameters to be equal. Model 7 is a limiting case of Model 8 in the same way as Model 2 is a limiting case of Model 3.

### 2.1 Parameter Values

In our study, we investigated nine distinct models to understand the variability in parameter values used to represent key phenotypic characteristics of TCR-pMHC interactions, such as specificity, sensitivity, and antigen discrimination. We observed considerable differences in parameter values reported across the literature, underscoring the challenge of selecting consistent values for accurate model comparison. To establish a reliable set of parameters, we drew data from four primary sources [28, 29, 33, 34], each contributing valuable insights but with varying emphasis on parameter ranges. Given these differences, we chose three representative values—mean, maximum, and minimum—for each parameter in our simulations. This approach allowed us to capture a comprehensive range of possible outcomes and enhanced the robustness of our model comparisons. Furthermore, to minimize dependency on individual parameters and ensure comparability, we applied a common parameter set across all models. This standard set is provided in the table below, with parameters unique to each model detailed in the subsequent sections.

- In model (3), KPR with limited signaling [28], the value *ξ* = 0.09 is used, similarly for (8), (9).
- In model (4), KPR with sustained signaling [28], the value Ω = 0.001 is used, similarly for (9).
- In model (6), KPR with activation chain stabilisation [30], *r* = 1.5 in equation for *k*_off_(*i*), and *r* = 1.03 in equation for *k*_*p*_(*i*), [Equations 8, 11 in [30]].
- In model (8), KPR with limited signaling and incoherent feed forward loop [31], *d* = 500*s*^−1^; *c* = 1*s*^−1^; *b* = 500*s*^−1^; *a* = 1*s*^−1^; *m* = 100; *l* = 100; *δ* = 50*s*^−1^; *µ* = 2.5*s*^−1^; *σ* = 0.5*s*^−1^.

### 2.2 Nature of solutions

All the mathematical models considered in what follows are systems of ordinary differential equations (ODE) obtained from reaction networks by assuming mass action kinetics. The equations are of the form 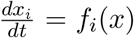 where the *x*_*i*_, 1 ≤ *i* ≤ *n*, are concentrations of certain substances and *x* is the vector with components *x*_*i*_. The functions *f*_*i*_ are polynomials and so, in particular, Lipschitz continuous. Thus for any time *t*_0_ and initial concentrations 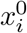 there exists a unique solution *x*_*i*_(*t*) on some time interval [*t*_0_, *t*_1_) with 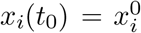. Proofs of this and other standard results on ODE used in what follows can be found in [35]. Because of their biological interpretation it is assumed that 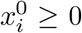 for all *i* and when this assumption is made it follows that *x*_*i*_(*t*) ≥ 0 for all *t* ≥ *t*_0_. To prove this note that (see for instance [36], Lemma 1) if 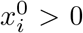 for all *i* it follows that *x*_*i*_(*t*) *>* 0 for all *t ≥ t*_0_. The more general statement then follows from the fact that solutions depend continuously on their initial data. As a consequence the non-negative orthant, i.e. the region defined by the inequalities *x*_*i*_ ≥ 0, is invariant. These reaction networks have conserved quantities *S*_*i*_ which are linear functions of the unknowns with non-negative coefficients satisfying 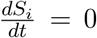. Thus *S*_*i*_ is equal to its value 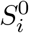 at *t* = *t*_0_. If there are *k* conserved quantities then by assigning them positive values 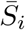 they may be used to eliminate *k* of the unknowns to obtain a reduced system. By ordering the unknowns suitably we can assume the variables eliminated are the *x*_*i*_ with *n* − *k* + 1 ≤ *i* ≤ *n* and we obtain a closed system of equations for the unknowns *x*_*i*_ with 1 ≤ *i* ≤ *n* − *k*. It is defined on a subset *K* of the non-negative orthant in ℝ^*n*−*k*^ given by the conditions that the original unknowns *x*_*i*_ are non-negative. *K* is an invariant set for solutions of the system. In all the examples considered in what follows each variable *x*_*i*_ occurs with a non-zero coefficient in at least one of the conserved quantities. It follows that *K* is compact. This implies that the solutions exist globally, i.e. on the interval [*t*_0_, *∞*). In fact in all these models there are exactly two conserved quantities, the total amount of ligand *L*_*T*_ and the total amount of receptors *R*_*T*_.

We will consider solutions where *x*_*i*_ *>* 0 for all *i*. In other words we restrict the system to the positive orthant. The corresponding region for the reduced system, the interior of *K*, will be referred to as the biologically relevant region. It turns out that for all the systems we consider if a solution lies in the biologically relevant region all its *ω*-limit points are also in that region. Proving this requires detailed consideration of the individual models but there is one part of the proof which applies to all the models and may be summed up in the following lemma.

#### Lemma 2.1.

*Let* 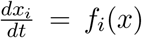 *be a system of ordinary differential equations arising from a reaction network by assuming mass action kinetics. It is always the case that f*_*i*_(*x*) = − *x*_*i*_*g*_*i*_(*x*) + *h*_*i*_(*x*) *for polynomials g*_*i*_ *and h*_*i*_. *Let x*(*t*) *be a solution in the non-negative orthant. Suppose that if at any ω-limit point of this solution the condition x*_*i*_ = 0 *for some i implies that h*_*i*_(*x*) *>* 0. *Then every ω-limit point of the original solution lies in the positive orthant*.

*Proof*. Suppose there exists an *ω*-limit point *x*^∗^ of the solution *x*(*t*) and some *j* for which 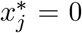. Let *y*_*i*_(*t*) be the solution with 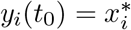 for all *i*. Then *y*(*t*) exists on the interval (−∞, ∞) and lies in the *ω*-limit set of *x*(*t*) and, in particular, in the non-negative orthant. Hence *y*_*i*_(*t*) ≥ 0 for all *t* ∈ (−∞, ∞) and all *i*. Since *y*_*j*_(*t*_0_) = 0 it follows that 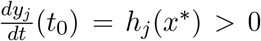. This implies that *y*_*j*_(*t*) *<* 0 for *t* slightly less than *t*_0_, a contradiction. ◼

Note that there is a correspondence between points on the boundary of *K* and points of the boundary of the non-negative orthant in ℝ^*n*^. Thus if Lemma 1 applies to the original system it can be concluded that any *ω*-limit point of a solution in *K* of the reduced system is in the interior of *K*.

We investigated the late-time behavior of our models, focusing on the existence and uniqueness of steady states and the convergence properties of solutions as *t* → ∞. Specifically, we address the question: Does each choice of parameters yield a unique steady state, and do all solutions converge to this steady state in the long term? Our findings are as follows:

1. For the Occupancy Model, the one-dimensional nature of system ensures convergence to a unique steady state.
2. For the KPR model and KPR with a stabilizing activation chain, convergence to a unique steady state is obtained.
3. For KPR with limited signaling, sustained signaling, and both, we establish convergence using the Deficiency Zero Theorem, as detailed below.
4. For KPR with negative feedback, the steady state is not always unique.
5. For KPR with IFF and limited signaling, convergence to a unique steady state is confirmed, as proven in subsequent sections.

The proof for KPR with limited signaling will now be written out as an example. The proofs for the other models are given in the appendix, section B

#### KPR with Limited Signaling

This model extends the kinetic proofreading concept, suggesting that once a TCR reaches the signaling-competent state *C*_*N*_, the bound TCR shifts to a non-signaling state *C*_*N*+1_ at a rate of *ξ*.

#### Mathematical formulation

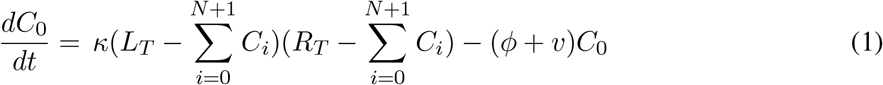

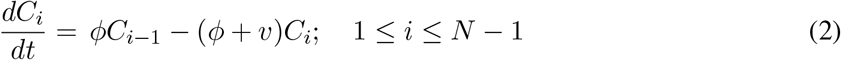

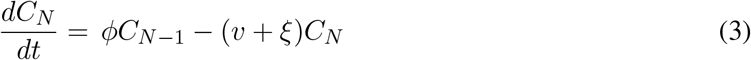

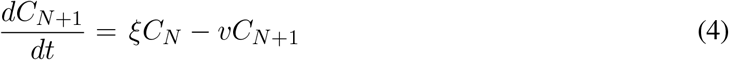

where *R*_*T*_ and *L*_*T*_ are the total concentrations of receptor and the ligand. If 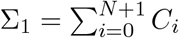 then equations (1)-(4) imply

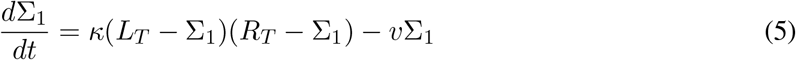

##### Lemma 2.2.

*Let* (*C*_0_(*t*), …, *C*_*N*+1_(*t*)) *be a solution of (1)-(4) contained in the closure K of the biologically relevant region. Then any ω-limit point* 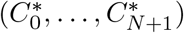 *of this solution is contained in the interior of K. In particular, any steady state is contained in the interior of K*.

*Proof*. The proof consists of repeated applications of Lemma 2.1. It follows from equation (5) that 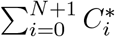 is strictly less than *L*_*T*_ and *R*_*T*_. It then follows from (1) that 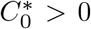. This in turn implies inductively, using (1)-(4) that 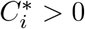. ◼

#### Stability of the solutions

For most of the models considered in what follows it will be shown using the Deficiency Zero Theorem that there is a unique steady state for each fixed choice of the conserved quantities and that this steady state is globally asymptotically stable. In general the deficiency of a chemical reaction network is given by *δ* = *n* − *l* − *s* where *n* denotes the number of species, *l* is the number of linkage classes and *s* denotes the rank of the reaction network. A linkage class is a connected component of the reaction graph. The rank of a reaction network is the number of elements in the largest set of linearly independent reaction vectors associated with that network [37]. It can be proved that the deficiency always satisfies the condition *δ* ≥ 0. The Deficiency Zero Theorem [38] says that if a network has deficiency zero and possesses a property called weak reversibility then the corresponding system of ODE with mass action kinetics has a unique positive steady state in each stoichiometric compatibility class and that this steady state is asymptotically stable within its class. Mass action kinetics refers to a specific form of rate equation commonly used in chemical kinetics, where the reaction rate is directly proportional to the product of the concentrations of the reactants [39]. In the models considered in what follows the stoichiometric compatibility classes are the sets defined by fixed values of the conserved quantities. In the case that no solution has an *ω*-limit point on the boundary the steady state is even globally asymptotically stable in its class. This means that any solution in the class converges to the unique steady state in that class.

In the case of KPR with limited signaling, the binding reaction has the form *L* + *R* → *C*_0_, where the bound receptor complex *C*_0_ undergoes sequential phosphorylation steps until it reaches the signaling competent state *C*_*N*_. After achieving this state, the bound TCR shifts into a state where it no longer signals *C*_*N*+1_ at a rate *ξ*. Each intermediate state *C*_*i*_ can decay, releasing *L, R*, and phosphate groups, making the network weakly reversible.

In our system, we have *n* = *N* +3 complexes, with a single linkage class (i.e., *l* = 1). To demonstrate that the deficiency of this network is zero, it suffices to show that the rank of the system is *s* = *N* + 2.

The *N* + 3 complexes are {*L* + *R, C*_0_, *C*_1_, *C*_2_, …, *C*_*N*_, *C*_*N*+1_}.

The stoichiometric matrix is represented as:

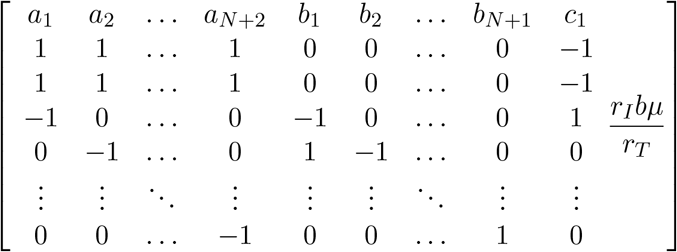

The first *N* + 2 columns of this matrix are linearly independent. Hence for this model *s* ≥ *N* + 2 and *δ* ≤ (*N* + 3) − (*N* + 2) − 1 = 0. Since *δ* is always non-negative this implies that *δ* = 0.

Given that this reaction network has a deficiency of zero and is weakly reversible, we can utilize the zero deficiency theorem. It follows that for the corresponding dynamical system with mass action kinetics there is exactly one steady state in each stoichiometric compatibility class and, as a consequence of Lemma 2.2, the steady state is globally asymptotically stable within its class.

### 2.3 Quantitative predictions of the models

We calculated mathematically the equations for response, maximum efficiency (*E*_max_), and potency (*EC*_50_) for all the models considered. In the following equations, *R*_*T*_ is the total amount of TCR receptor and *L*_*T*_ is the total amount of ligand. Also 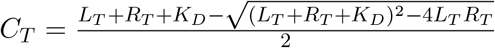, where *K*_*D*_ = *k*_off_ */k*_on_.

In all equations 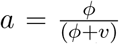. For other models the expressions for *r*_*±*_, *δ, ϵ, W, U*, etc. can be found in the appendix. Analytic observations on monotonicity and non-monotonicity of response functions in our model plots has been discussed in detail in the appendix.

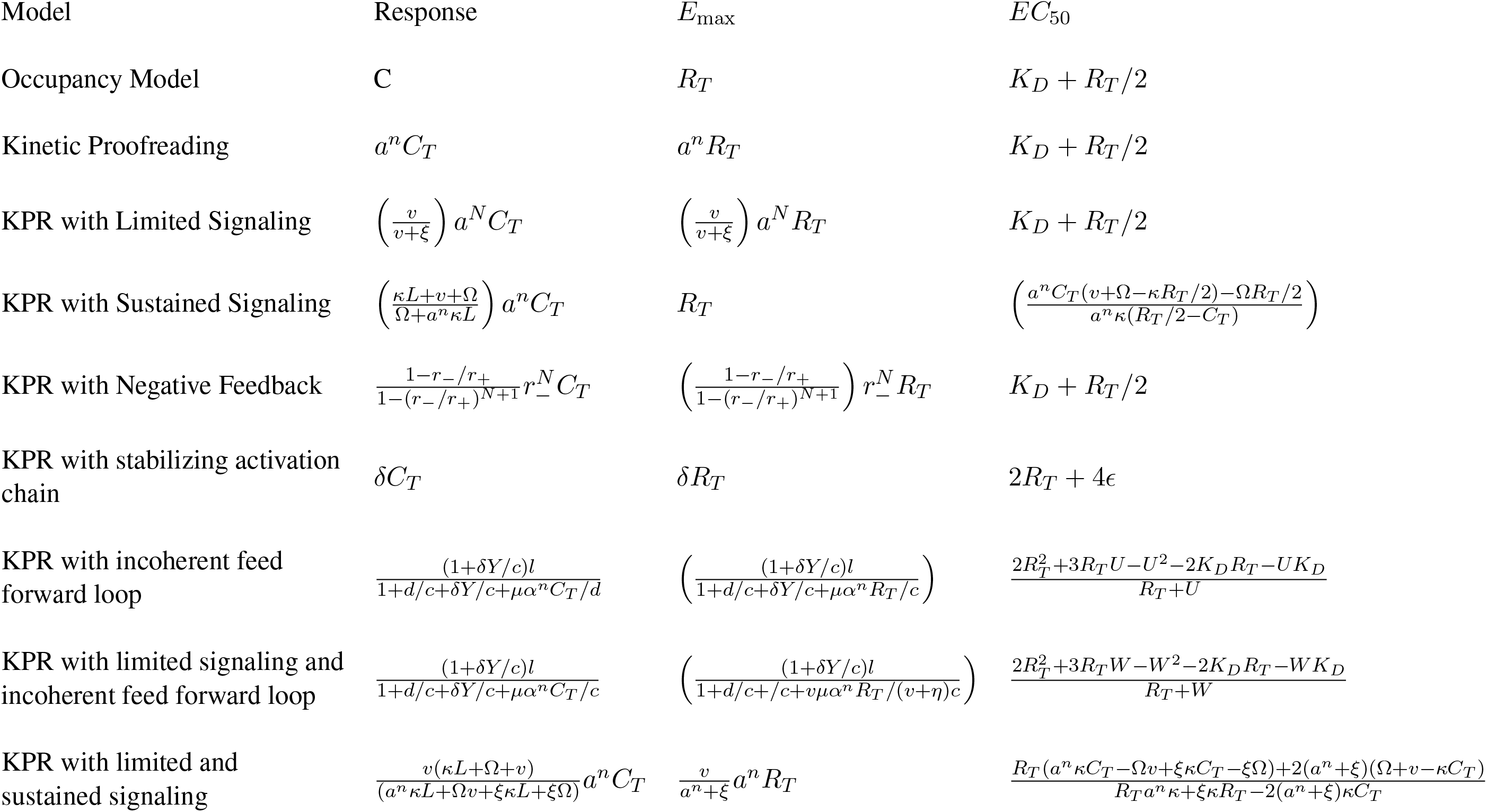

**Fig. 1.**
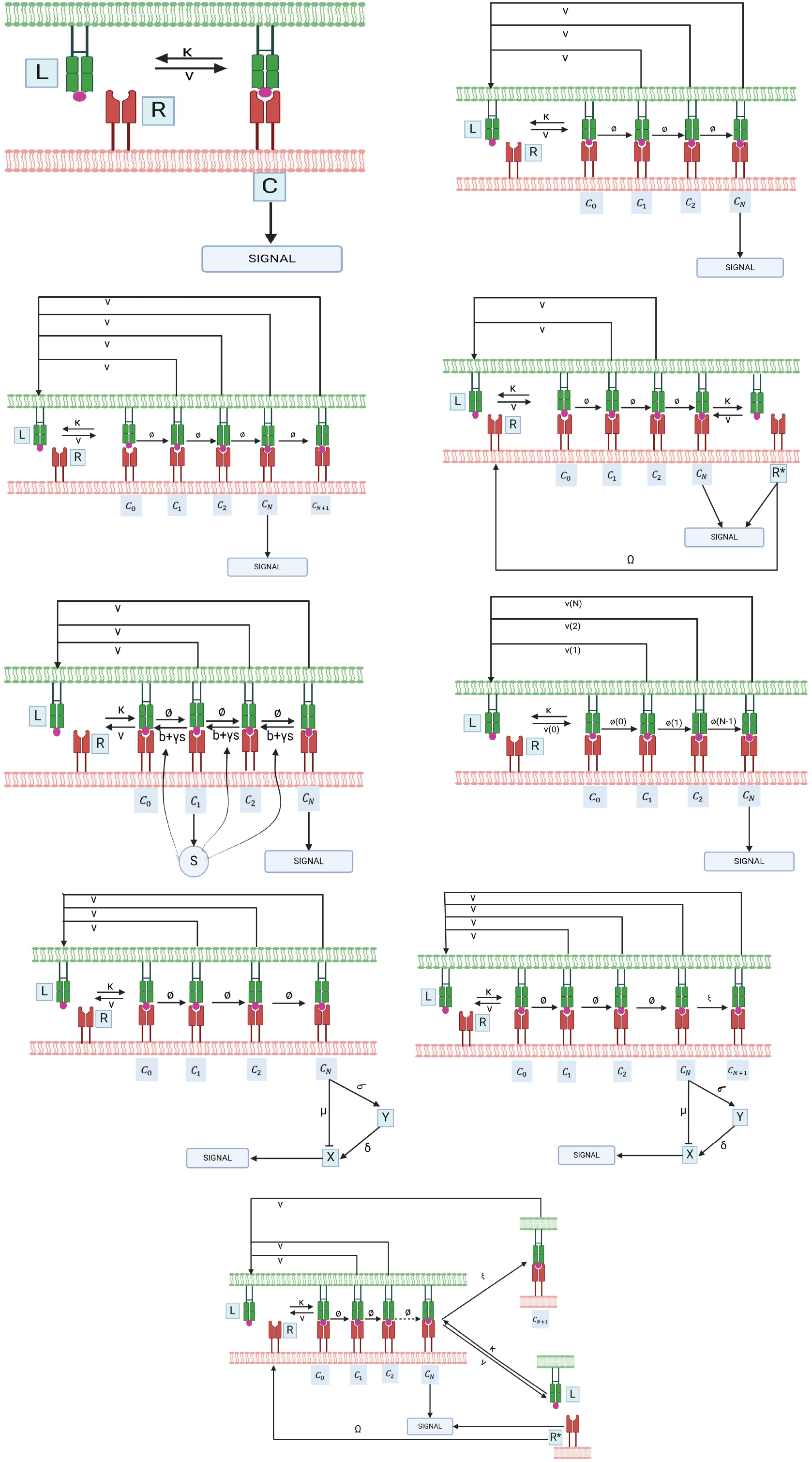
Phenotypic models of T cell activation

### 2.4 Numerical simulations

**Fig. 2.**
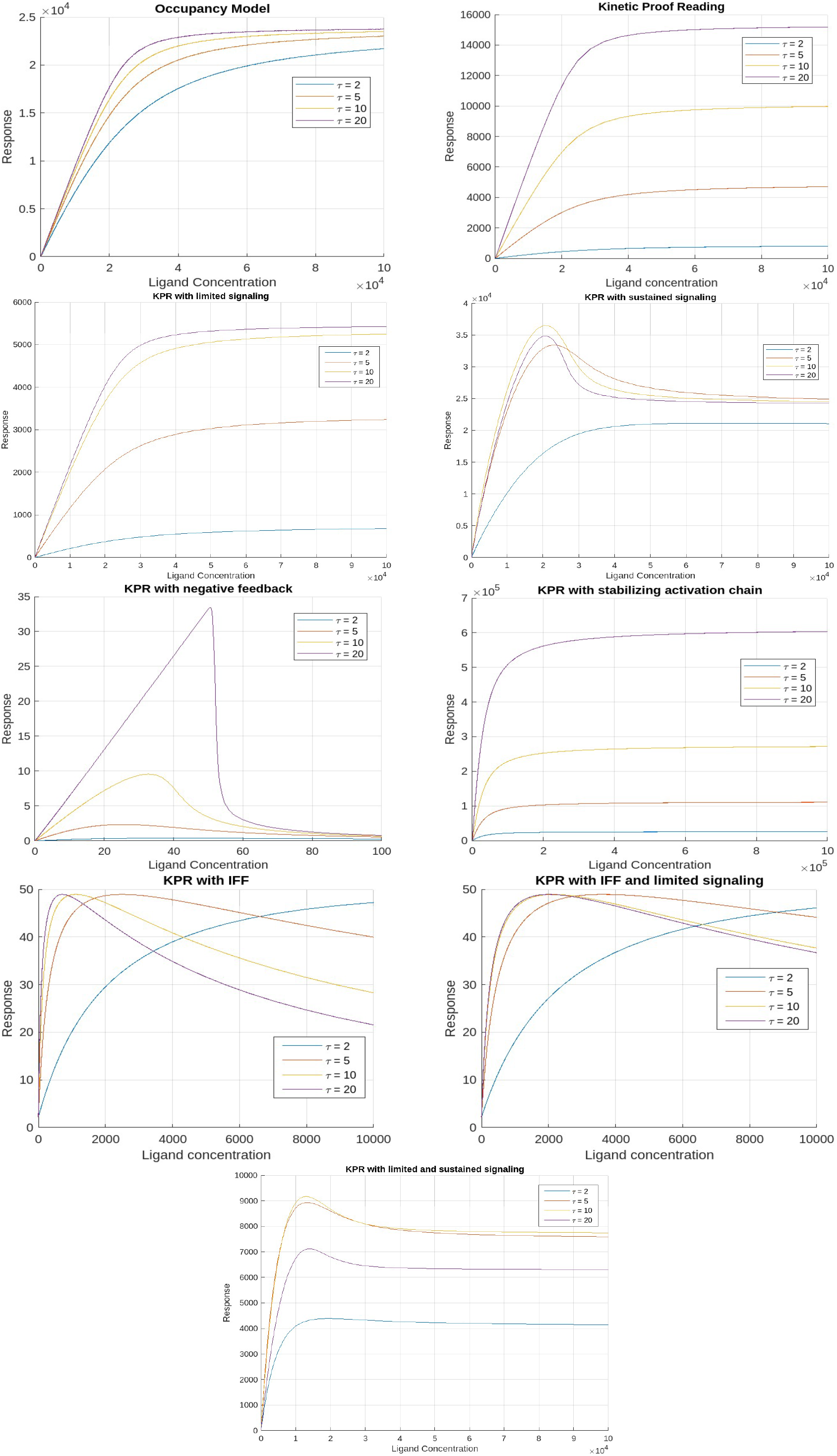
Predicted dose-response patterns for ligands with progressively longer dissociation times (*τ*) (based on mean parameter values).

**Fig. 3.**
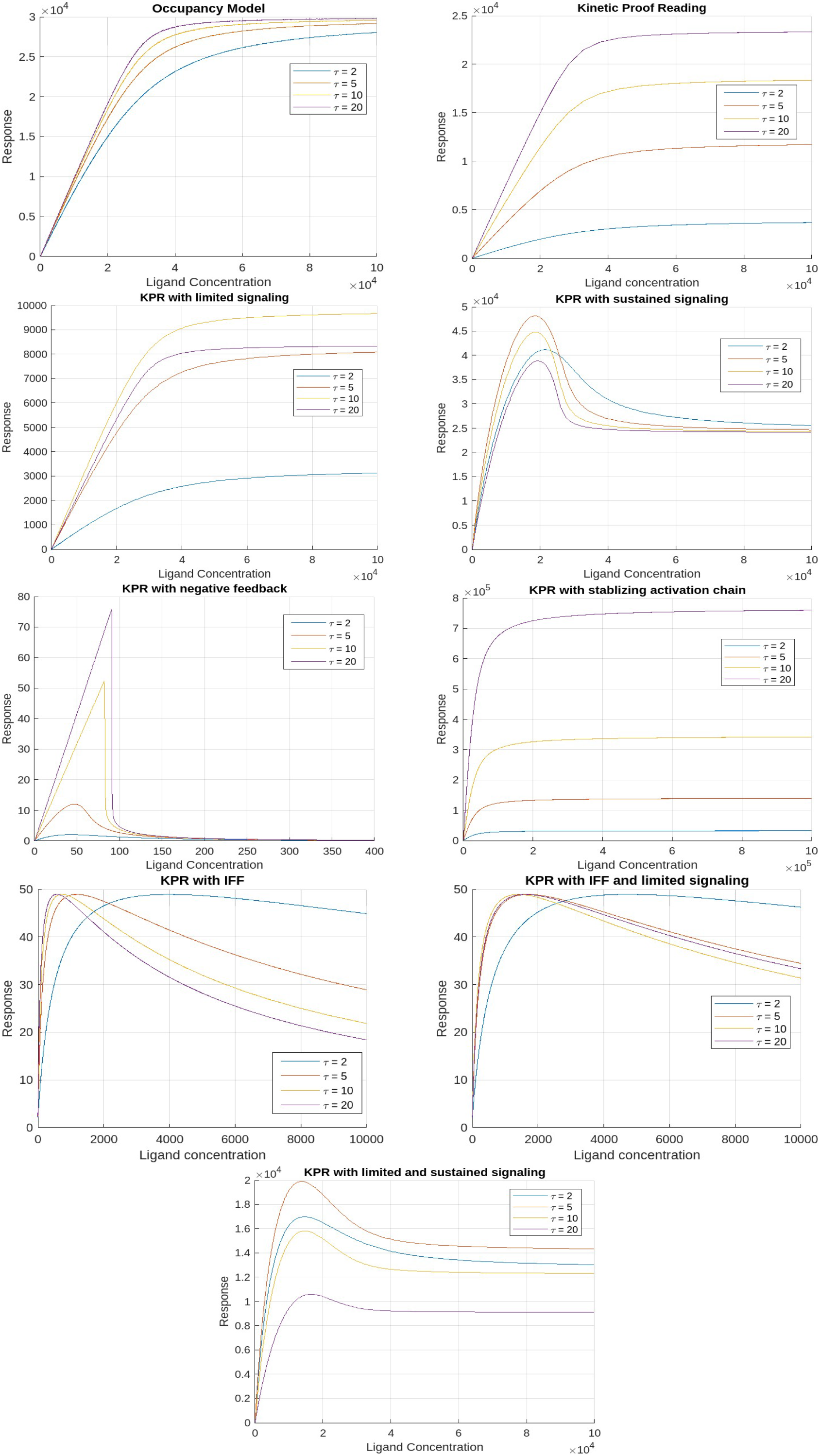
Predicted dose-response patterns for ligands with progressively longer dissociation times (*τ*) (based on maximum parameter values).

**Fig. 4.**
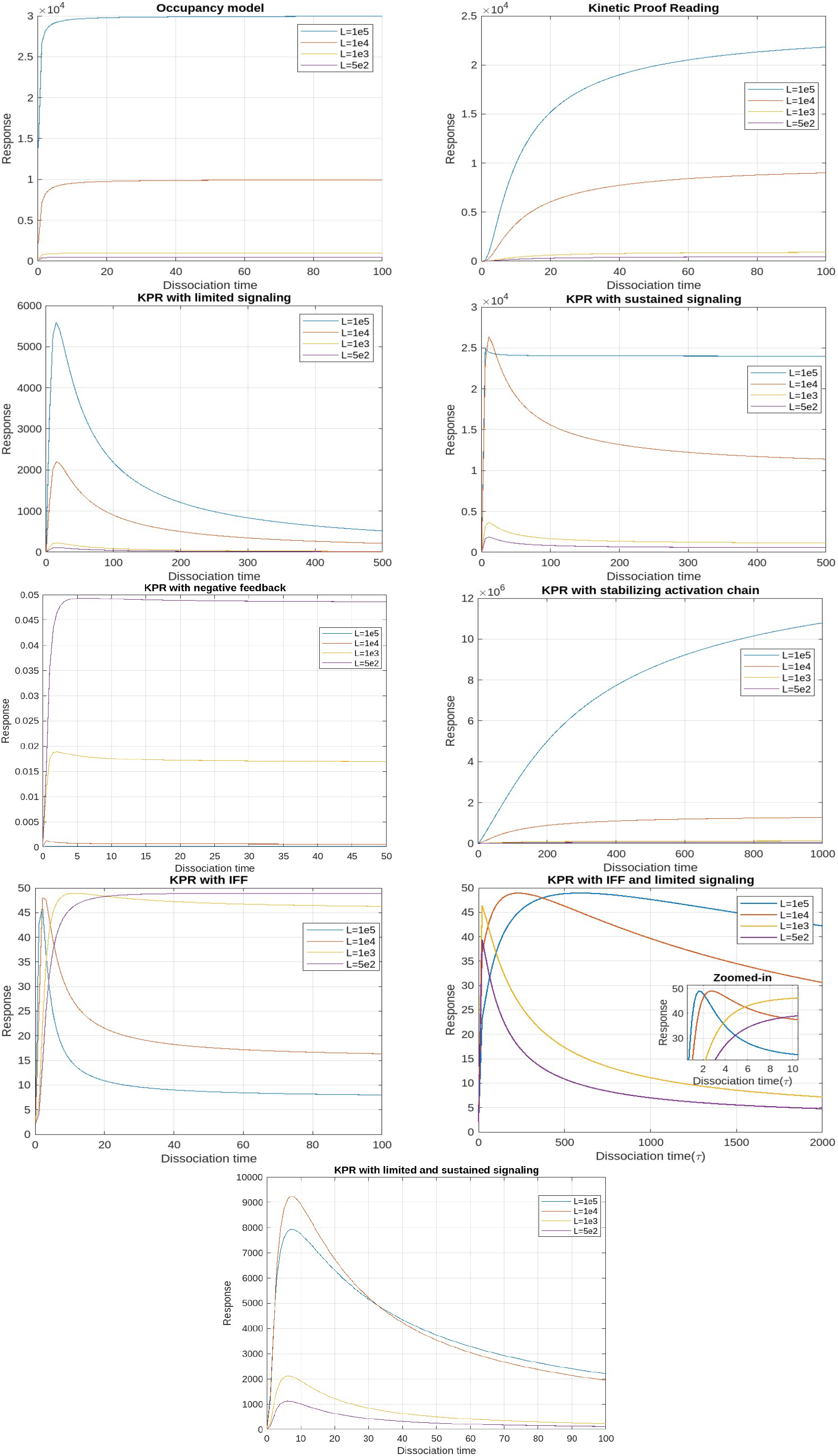
T cell activation as a function of ligand dissociation time (*τ*) at a constant ligand concentration (based on mean parameter values).

**Fig. 5.**
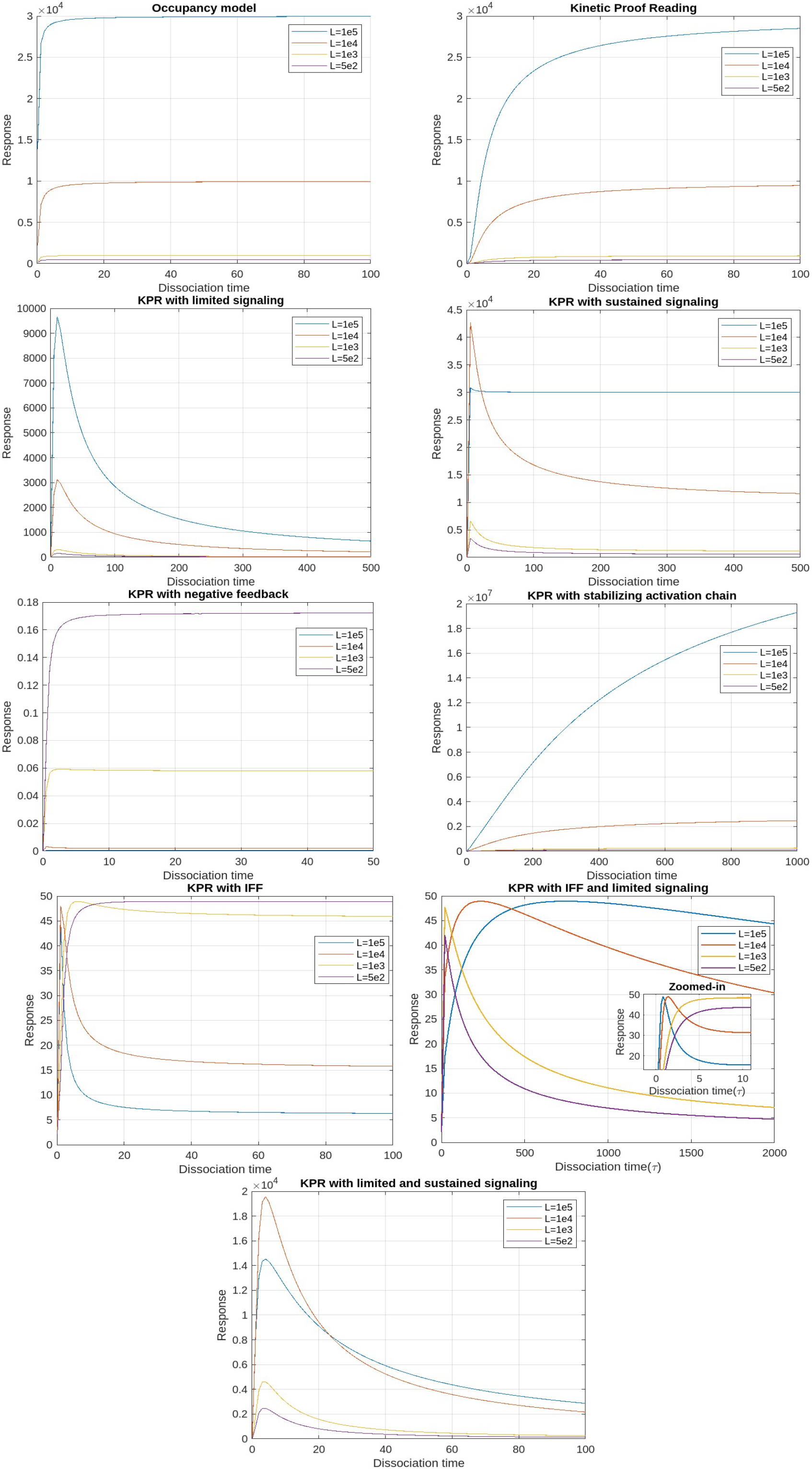
T cell activation as a function of ligand dissociation time (*τ*) at a fixed ligand dose (based on maximum values of parameters)

**Fig. 6.**
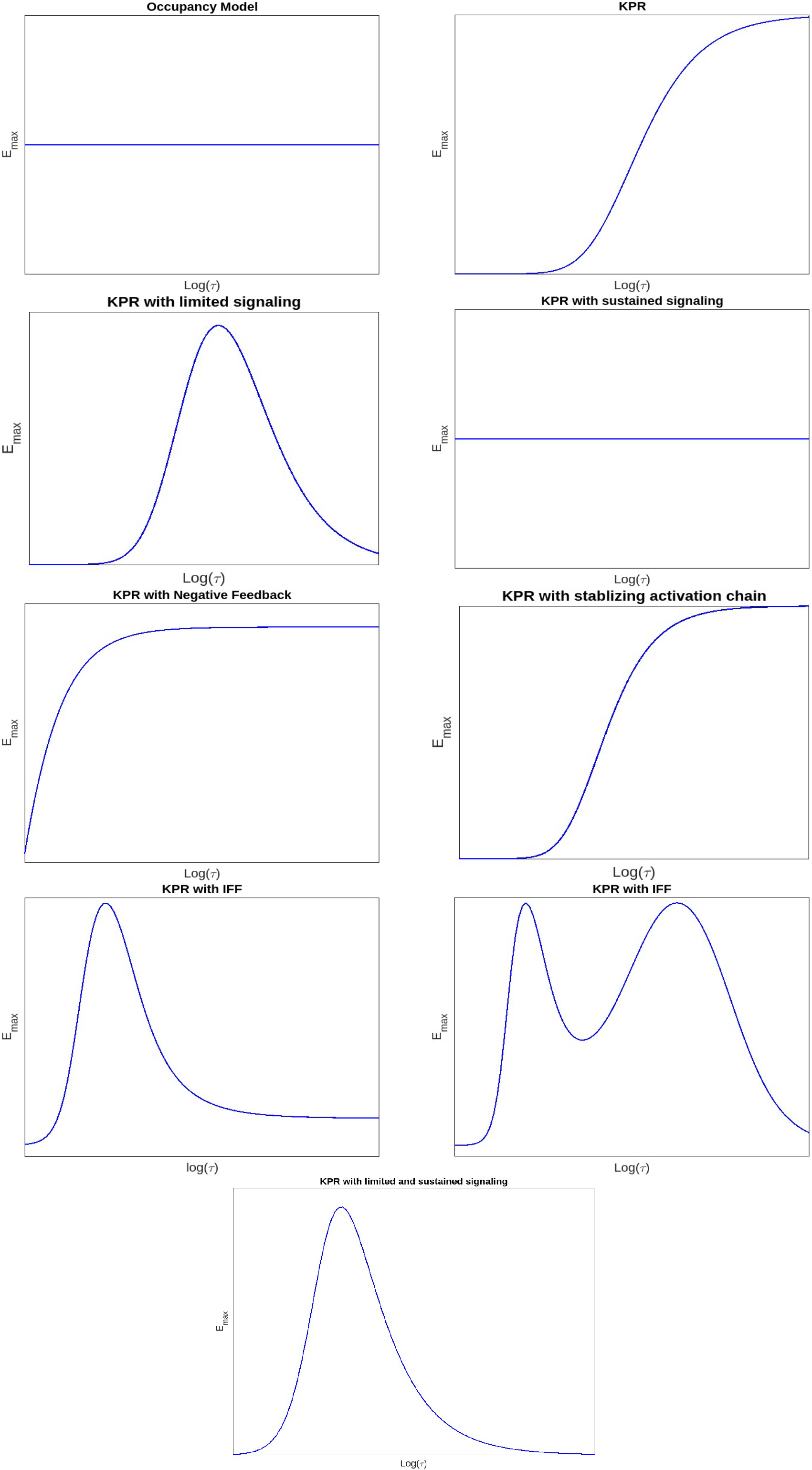
Plot of *E*_max_ vs dissociation time *τ*

**Fig. 7.**
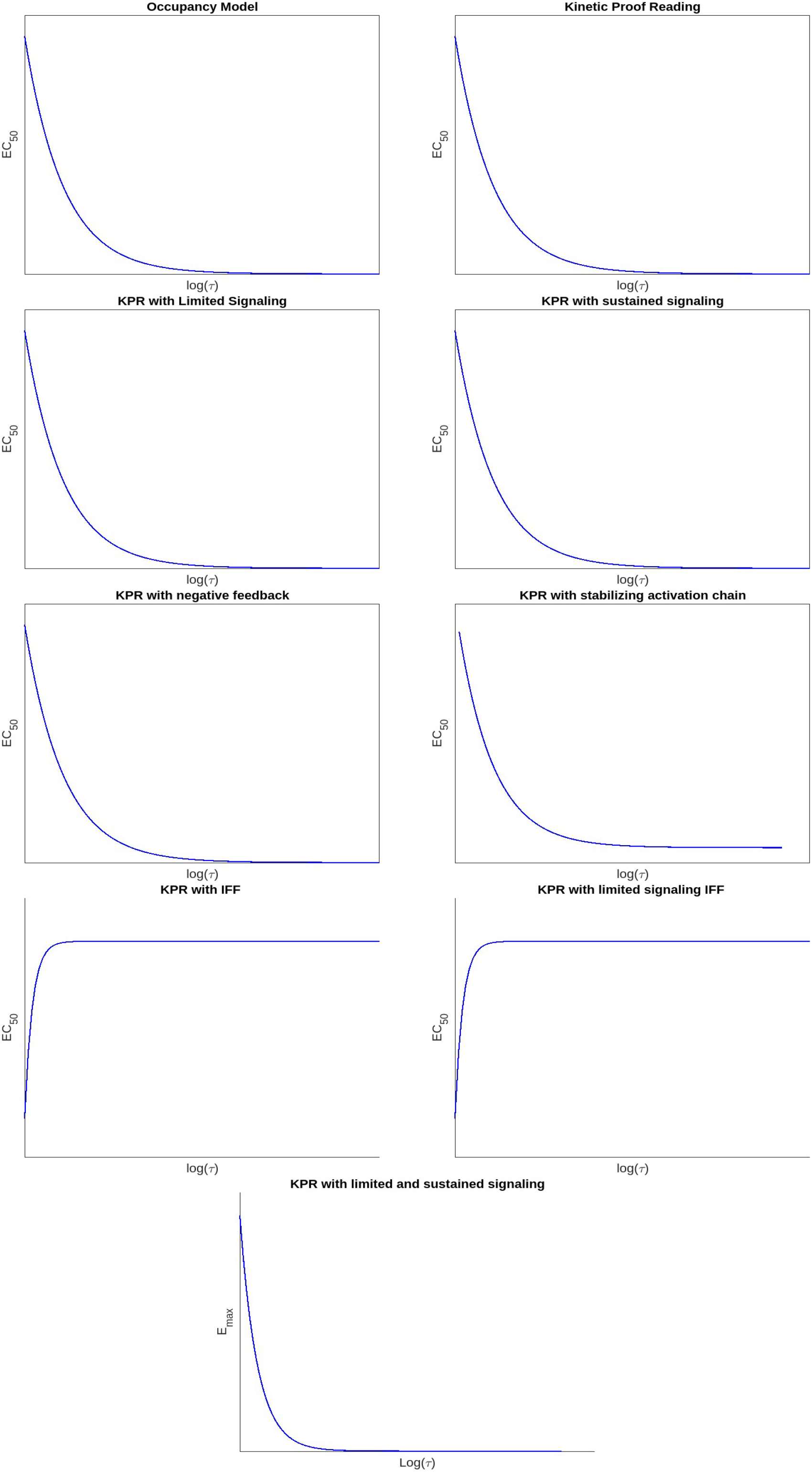
Plot of *EC*_50_ vs dissociation time *τ*

## 3 Model Characteristics and Results

A phenotypic model is a conceptual or mathematical framework for describing observable traits (phenotypes) in a biological system. With minimal assumptions, these models replicate experimental data, offering advantages such as fewer unknown parameters and easier interpretation. Unlike mechanistic models, phenotypic models make no explicit assumptions about TCR triggering mechanisms, allowing compatibility with all known TCR activation processes. For T cell activation, key experimental factors include TCR-pMHC engagement duration, bond affinity, ligand threshold concentration, and optimal dwell time.

Our analysis yielded several novel insights. In the sustained signaling model, we found an optimal response with respect to ligand concentration for all dissociation time values. In the negative signaling model, we observed an optimal response with respect to dissociation time. While this optimum is not very pronounced, it agrees with predictions made in [25]. For IFF, we found that the dose-response curves intersect at different ligand concentrations and dissociation times. Moreover, we discovered that ligands with longer dissociation times generate the strongest response before reaching the peak contradicting the conclusions made earlier [31]. For IFF with limited signaling, we found unique bimodal response patterns with respect to dissociation time.

In this section, a concise overview is presented, covering the phenotypic models employed in this study, their experimental backing, mathematical assumptions, and the degrees of intricacy. The mathematical formulations and analysis of these models are present in appendix A

### Occupancy model

This model serves as the foundation for explaining the binding of TCRs to pMHC and the subsequent T cell response. It posits that the activation of T cells is directly connected to the number of TCRs which are engaged at equilibrium. When a pMHC binds to a TCR, the TCR instantly transitions into a state capable of signaling. For this model, we examined the relationship between T cell response as a function of ligand concentration, and dissociation time. The results revealed that, within the occupancy model, there is no optimal response level. Also from the plots it is evident that, the maximum response, *E*_max_, remains constant regardless of parameter variations, because *E*_max_ is independent of binding concentration, any pMHC, when present in sufficiently high concentrations, can elicit the maximum T-cell response. Consequently, even low-affinity pMHCs can trigger a response if their concentration is increased sufficiently, indicating that this model fails to account for antigen discrimination. Hence this model fails to reproduce experimentally observed characteristics of T cell response.

### Kinetic Proofreading

This model proposes that for the binding of TCR-pMHC to effectively trigger a signaling response, the interaction must persist long enough for the TCR to reach a signaling-competent state. For this framework, a pMHC ligand (L) can reversibly bind to a TCR receptor (R) to form a pMHC-TCR complex, denoted as *C*_0_. Upon binding, this complex undergoes a series of biochemical modifications, ultimately reaching a signaling-competent state *C*_*N*_. If the pMHC unbinds from the TCR at any intermediate stage, these modifications are immediately reversed, returning the TCR to its unmodified state. Thus in this model, T cell activation is determined by the proportion of TCRs in the *C*_*N*_ state, rather than the overall number of TCRs that are occupied. This suggests that successful activation relies on the fraction of TCRs that remain bound to pMHC complexes long enough to reach the *C*_*N*_ state.

These biochemical modifications required to reach a signalling state encompass various mechanical processes involved in T cell activation, like initial dimerization of receptors, the phosphorylation of multiple tyrosine residues within the TCR complex, and the subsequent mobilisation and stimulation of ZAP-70. In fact, experimental findings have revealed differences in the early phosphorylation patterns of molecules like p21/p23 and in the mobilisation and activity of ZAP-70 in return to pMHCs of varying potencies [8, 9].

The plots demonstrate that the relationship between response and dissociation time does not show an optimal point; however, the maximum response (*E*_max_) is dependent on pMHC dissociation time. This suggests that merely increasing the concentration of pMHCs with shorter dissociation times will not result in the same level of response as pMHCs with longer dissociation times [40, 6]. Thus, the KPR model elucidates the mechanism of antigen discrimination based on dissociation time. Unlike the occupancy model, where pMHC concentration and affinity can compensate for each other, the KPR model shows that high pMHC concentrations cannot offset a short dissociation time, leading to diminished T cell responses.

### Kinetic proofreading with Limited signaling

This model builds on the KPR concept and proposes that once a TCR achieves a signaling competent state *C*_*N*_, it transitions to a non-signaling state *C*_*N*+1_ at a rate *ξ*.

Unlike self pMHCs, antigenic pMHCs on the surface of APC are relatively sparse (tens to hundreds of copies among approximately 10^5^ MHC class I molecules for each APC [10]). This limited availability constrains the number of T cell receptors (TCRs) that can simultaneously bind to agonist pMHCs. Studies involving dual receptor T cell clones expressing two distinct *αβ* dimers have demonstrated that TCR activation is not transmitted to adjacent receptors, suggesting that each TCR must independently engage with pMHC to undergo activation.

To overcome this limitation, the serial triggering model proposes that a single antigenic pMHC can sequentially activate multiple TCRs over time [11]. This indicates that each TCR produces a single signaling packet for every instance of pMHC binding, which limits ongoing signaling from pMHC complexes that have longer dissociation times. Essentially, signaling through individual TCRs is limited. From a mechanistic standpoint, this is backed by findings that TCR signaling is limited to their transition from the outer edges to the center of the immunological synapse, and it stops once they are marked for eradication from the T cell surface [12, 41, 42].

The plots of the response versus dissociation time reveal an optimal dissociation time for the mean and maximum parameter values across all pMHC doses, although this optimal time is less apparent at the minimum values. This finding is consistent with experimental observations showing an optimal dissociation rate in both in vivo [43] and in vitro studies [44]. A possible explanation for this optimality, particularly at high pMHC doses, is that pMHC molecules with prolonged dissociation times may form stable yet unproductive complexes with TCRs, preventing them from sequentially engaging with other receptors. As a result, the most effective pMHC molecules are those that persist just long enough to activate TCRs for signaling but dissociate swiftly once signaling is complete. Notably, no optimal dissociation time is observed concerning ligand concentration.

### Kinetic proofreading with Sustained Signaling

This model is a further modification of the KPR model. According to this model, TCRs in the signaling-competent state *C*_*N*_, persist in signaling for a certain duration after the unbinding of pMHC. Subsequently, they revert to the baseline state at a rate Ω. Rather than assuming that the signal is restricted, this model proposes that even after the pMHC has detached, the signaling competent TCRs can sustain signaling for a specified duration. From a mechanistic perspective, this ties into the idea that TCRs and their associated complexes, known as signalosomes, keep transmitting signals even after pMHC has dissociated. This signaling remains active until phosphatases deactivate the TCRs through dephosphorylation or the TCRs are internalized.

The serial triggering model posits that rapid pMHC dissociation from previously engaged TCRs is crucial for enabling sequential receptor activation, which may appear at odds with the prolonged bond lifetimes suggested by the kinetic proofreading model. To bridge this gap, the concept of an optimal dwell time was introduced [13], proposing that intermediate half-lives are critical for effective downstream signaling. Experimental support for this idea comes mainly from studies using mutated TCR panels [14].

In addition, it was experimentally found that at low concentrations of pMHC there is an optimal dissociation time for T cell activation, but at high pMHC concentration there is no such requirement [14, 45]. The response plots with respect to dissociation time also reveal that there is an optimal dissociation time at lower doses. However, this optimal point disappears when doses are higher. This phenomenon can be explained by noticing that, at low doses, there is a delicate interplay among kinetic proofreading and serial binding. Consequently, the highest level of activation is achieved when the dissociation time is at an intermediate value. In contrast, at high pMHC doses, the need for pMHC to sequentially bind to TCRs to sustain a continuous signal diminishes. At higher pMHC concentrations, it becomes possible to produce a significant number of TCRs in a signaling-capable state, even for pMHCs that have rapid separation rates. While pMHCs with short dissociation times may have a lower probability of producing competent-signaling TCRs owing to kinetic proofreading, they have a unique ability. They can maintain a considerable population of signalling-competent TCRs at higher pMHC doses. This is achieved by binding to a sustained signaling TCR and preventing it from reverting to its baseline state. Consequently, the parameter *E*_max_, representing the maximum response, becomes independent of the dissociation time under these conditions.

The plots showing the response versus ligand concentration for both the mean and maximum values of the parameters also indicated an optimum. This observation is novel, as previously, an optimum with respect to ligand concentration had only been reported for the negative feedback model. Unlike the KPR model with negative feedback, the nature of this curve is that it initially rises, reaches a peak, and then levels off.

### Kinetic proofreading with negative feedback

The Kinetic Proofreading with Negative Feedback (KPR-NF) model refines the KPR framework by introducing the concept that the rates of complex formation within the activation chain can be modulated at various transitional states and/or within the last signaling state *C*_*N*_. This modulation is crucial for preventing excessive activation of T cells and plays a significant role in determining the sensitivity and duration of T cell activation. The modulation is facilitated through a single negative feedback mechanism mediated by enzymes like SHP-1 (Src homology region 2 domain-containing phosphatase-1), which disrupt the activation cascade, thereby regulating T cell activation.

Plots illustrating the response versus ligand concentration show that the response reaches an optimum for mean, maximum, and minimum parameter values before rapidly decaying to near zero beyond the peak for all time values. These bell-shaped curves can be explained by observing that, at low ligand concentrations, the T cell response rises as the ligand concentration increases. This increase continues until the negative regulator predominates, causing the response to decrease as the inhibitor’s quantity rises. This optimal response for pMHC dose has also been predicted by previous studies [28, 29]. However, an optimum concerning dissociation time was not achieved. It has been shown earlier that such an optimum with respect to dissociation time is achievable under certain parameter conditions[25]. Our plots confirm this prediction, showing that KPR with negative feedback curves presents an optimum response as a function of dissociation time for some ligand concentrations, although this optimum is not very pronounced. The plot of *E*_max_ versus dissociation time indicates that the response is dependent on pMHC dissociation times.

### Kinetic proofreading with stabilizing activation chain

This model is based on the premise that the proofreading mechanism operates differently for non-self and self-pMHC peptides. Consequently, the activation chain for these ligands exhibits contrasting properties. The model suggests that KPR complexes stabilize foreign peptides while weakening their interaction with self-peptides. As the activation chain progresses, this differential behavior results in an enhanced response, characterized by increased sensitivity and specificity [30]. As the proofreading process continues, strengthening or weakening of the *C*_*i*_ complexes, along with fluctuations in activation time, is represented by adjustments in the rate constants *v*(*i*) (for *i* = 0, 1, …, *N*) and *ϕ*(*i*) (for *i* = 0, 1, …, *N* − 1).

Despite the assumption that this model refines the selectivity and specificity of the T cell activation process, the plots tell a different story. Neither response as a function of dissociation time nor ligand concentration shows an optimum. However, plots for KPR with sustained signaling indicate that *E*_max_ depends on pMHC dissociation time. Therefore, pMHCs with shorter dissociation times are unable to generate the same level of response as those with longer dissociation times, even if their concentrations are increased. Therefore, the KPR model with a stabilizing activation chain demonstrates antigen differentiation determined by dissociation time but fails to exhibit any observable phenotypic characteristics.

### Kinetic proofreading with incoherent feed forward loop

This model was initially proposed in [31], integrating KPR with an IFF. The incoherent feed forward loop is a common pattern found in molecular regulatory networks, capable of generating biphasic responses either based on time or dosage. In mathematical terms, this motif involves a specific arrangement where a single input influences a single result by passing through multiple intermediate pathways. In this model, when the pMHC ligand binds to the TCR, it transitions to the signaling component state *C*_*N*_, which directly inhibits X. However, by activating Y, which in turn can activate X, *C*_*N*_ indirectly activates X as well. In the context of the proposed model, the incoherent feed-forward loop fine-tunes T cell activation. It ensures that only high-affinity interactions (those likely to be correct) are sustained long enough to pass through the kinetic proofreading steps, while lower-affinity interactions are rapidly inhibited.

The plots for response as a function of ligand concentration for mean and maximum values shows an optimum for higher dissociation time. However the optimum is lost at lower values. Also, the ligand with higher dissociation time produces the greatest response occurring left of the peak and this result is opposite to the result of [31], on the basis of which they rejected this model.

In addition, response with respect to dissociation time for mean and maximum values, shows an optimum for higher doses. However this optimum is not observed at lower doses. These results are opposite to the results shown by KPR with sustained signaling. This result can be explained by the fact that low-affinity pMHCs (peptide-MHC complexes) produce a lower peak level of *C*_*N*_ compared to high-affinity pMHCs. If this peak level is below the point where *Y* becomes saturated, then even at high concentrations, low-affinity pMHCs will not cause inhibition when they interact with *Y*.

As the model predicts an optimum for *E*_max_, it suggests that the dose-response curves it generates might intersect for certain dissociation times. This characteristic has been evident in plots for this model. While not explicitly stated, earlier studies suggest the potential for intersection in dose–response curves, highlighting that the effectiveness of antigens is affected not only by how well they bind but also by the particular dosage used during administration. [40, 23]. The combined model effectively accounts for a significant portion of the observed phenotype characteristics.

### Kinetic proofreading with limited signaling and incoherent feed forward loop

This model extends the KPR mechanism by incorporating limited signaling and an incoherent feed-forward loop. It assumes that TCR signaling is time-restricted: once a TCR binds to its pMHC complex, it signals only for a limited period. After this window, the bound TCR shifts to a non-signaling transit state. Specifically, active TCR–pMHC complexes (*C*_*N*_) can signal briefly before converting to a non-signaling state (*C*_*N*+1_). This conversion penalizes pMHC molecules with prolonged binding, reducing signaling efficacy and adding quality control to antigen recognition, with IFF this signal is further enhanced.

The response plots as a function of the concentration of the ligand for the mean and maximum values exhibit distinct patterns. High-affinity ligands demonstrate an optimal response at specific concentrations, whereas this optimum diminishes with lower affinities. Notably, high-affinity ligands generate the strongest response on the left side of the peak, consistent with findings from [31].

The response plots as a function of dissociation time reveal complex regulatory dynamics. For both mean and maximum values, we observed bimodal response patterns. This bimodal response was also evident in the plot of *E*_max_. While less frequent, bimodal responses have also been reported previously by other researchers. These bimodal responses describe sequential phases of activation, often observed in cytokine regulation and metabolic adaptation [46, 47]. For instance, IL-6 exhibits a bimodal role in immune responses, initially promoting T cell activation but later exerting immunosuppressive effects [48]. These dual-phase mechanisms play critical roles in immune homeostasis and have implications for therapeutic interventions, including checkpoint blockade therapy and metabolic reprogramming in T cells.

### Kinetic proofreading with limited and sustained signaling

This model has been formulated by combining two KPR models, (3) and (4). It integrates the concept that while a TCR is limited to generating a single signaling packet per pMHC binding event, the persistence of signaling after pMHC unbinding enhances the overall signaling duration. This extended signaling duration amplifies the T cell response, providing a more robust immune reaction.

The response shows an optimum as a function of dissociation time for both the mean and maximum values of the parameters considered. This result aligns with KPR with limited signaling, but the dose-response curves here are more prominent. Unlike models with sustained signaling where the optimum may be lost at high pMHC doses, this combined model retains the optimum even at high pMHC levels. Additionally, the response shows an optimum with ligand concentration, consistent with the KPR with sustained signaling. The *E*_max_ also exhibits an optimum, implying that the dose-response curves generated by this model could overlap at particular dissociation times. This property is reflected in the model plots, indicating that at low doses, one antigen may induce superior T cell activation compared to another, while at high doses, their relative performance may be reversed [40, 23].

In summary, by incorporating aspects of both limited and sustained signaling models, this combined approach provides a nuanced understanding of TCR signaling. It highlights how optimal dissociation times and ligand concentrations can fine-tune the immune response, ensuring effective T cell activation across different antigen doses.

#### Validation of Previous Findings

Two major findings discussed by [25] highlight the presence of three positive steady states for certain parameter sets and the impossibility of exceeding this number. Additionally, the response function exhibited non-monotonic behavior with respect to dissociation time under specific parameter values. While these results were intriguing and novel, the majority of the parameter values used in their study were arbitrarily chosen.

To investigate whether these findings hold true under experimentally and biologically plausible parameter sets, we extended the analysis by comparing the results using parameters derived from four distinct sources.

Our analysis confirmed the existence of exactly three positive steady states, all of which satisfied the biologically feasible region criteria. We also evaluated the stability of these steady states and found that two of them were stable. Furthermore, we observed that the response function exhibited an optimal behavior with respect to dissociation time. While the mean parameter values from our sources revealed a modest optimal response, a slight increase in the phosphorylation rate resulted in a more pronounced optimal response, as illustrated in the accompanying figure 8.

**Fig. 8.**
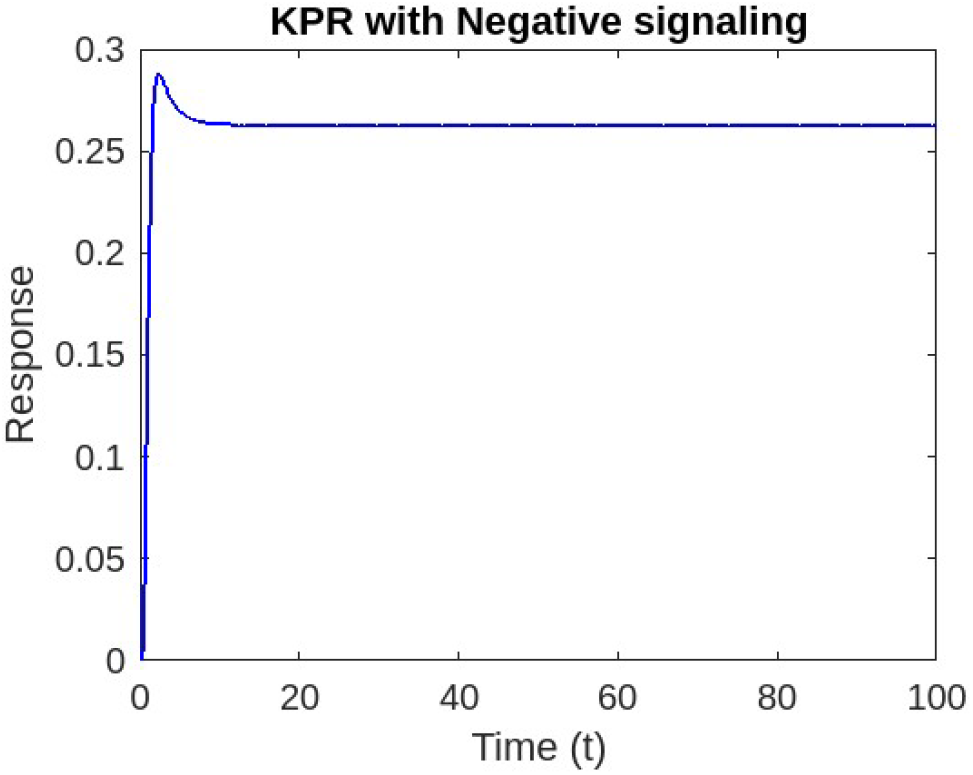
Response as function of Dissociation time at a fixed Ligand Concentraion

## 4 Sensitivity analysis

Sensitivity analysis is an essential tool for studying biological systems due to its ability to reveal how variations in parameters influence system behavior, highlighting the factors with the greatest impact on outputs. It also supports the validation and refinement of mathematical models by identifying parameters that need precise measurements versus those that can vary, enhancing model robustness. A highly effective method for examining parameter uncertainty is the Latin Hypercube Sampling-Partial Rank Correlation Coefficient (LHS-PRCC) approach. This technique efficiently explores the full parameter space with an optimal number of simulations, combining Latin Hypercube Sampling (LHS), introduced by McKay in 1979 [49], and Partial Rank Correlation Coefficient (PRCC) analysis [50]. LHS generates diverse parameter values within a defined range for each simulation, while PRCC assesses the relationships between these parameters and the model’s outputs [51]. PRCC, a sample-based technique, assesses the relationship between a model’s output variable and its parameters using sample points generated through Latin Hypercube Sampling (LHS). Sensitivity analysis with LHS-PRCC aims to identify the most influential parameters on model predictions and to rank these parameters by their impact on achieving accurate outcomes [52].

We conducted a sensitivity analysis using Latin Hypercube Sampling with Partial Rank Correlation Coefficients (LHS-PRCC) across all our model systems. Our objective was to identify which parameters significantly influence the model output. We began by treating the parameters as uncertain and subject to variation under Latin Hypercube Sampling (LHS). To evaluate their individual contributions to model predictions, we performed 10,000 simulations of the model. For each parameter, we considered both the highest and lowest values, with baseline values set to the mean. The minimum and maximum values were drawn from reported values in relevant literature, ensuring that each LHS parameter was based on established references in Table 1. A positive PRCC value indicates that an increase in the parameter leads to an increase in the model output. In contrast, a negative PRCC value means that as the parameter increases, the model output decreases.

**Table 1:** Mean, Maximum, and Minimum values derived from the sources for various parameters.

| Parameter | Source [29] | Source [28] | Source [33] | Source [34] |
| --- | --- | --- | --- | --- |
| Receptor per cell ($R_T$) | $3 \times 10^4$ | $1.5708 \times 10^4$ | $2 \times 10^4$ | $3 \times 10^4$ |
| Ligand/Receptor association rate ($\kappa$) | $10^{-4}$ | $3.1831 \times 10^{-5}$ | $10^{-4}$ | $10^{-5}$ |
| Phosphate per cell ($S_T$) | $6 \times 10^5$ | $6 \times 10^5$ | $6 \times 10^5$ | $3 \times 10^5$ |
| Phosphorylation rate ($\phi$) | 0.09 | 1 | 1 | 0.05 |
| Spontaneous dephosphorylation rate (b) | 0.04 | 0.04 | 0.04 | 0.02 |
| Phosphatase efficiency ($\gamma$) | $1.2 \times 10^{-6}$ | $4.4 \times 10^{-4}$ | $4.4 \times 10^{-4}$ | |
| SHP-1 activation rate ($\alpha$) | | | $2 \times 10^{-4}$ | $10^{-4}$ |
| SHP-1 deactivation rate ($\beta$) | | | 1 | $6 \times 10^{-3}$ |

| Parameter | Mean | Maximum | Minimum |
| --- | --- | --- | --- |
| Receptor per cell ($R_T$) | $2.4 \times 10^4$ | $3 \times 10^4$ | $1.5708 \times 10^4$ |
| Ligand/Receptor association rate ($\kappa$) | $6.05 \times 10^{-5}$ | $10^{-4}$ | $10^{-5}$ |
| Phosphate per cell ($S_T$) | $5.25 \times 10^5$ | $6 \times 10^5$ | $3 \times 10^5$ |
| Phosphorylation rate ($\phi$) | 0.535 | 1 | 0.05 |
| Spontaneous dephosphorylation rate (b) | 0.035 | 0.04 | 0.02 |
| Phosphatase efficiency ($\gamma$) | $2.93 \times 10^{-4}$ | $4.4 \times 10^{-4}$ | $1.2 \times 10^{-6}$ |
| SHP-1 activation rate ($\alpha$) | $1.5 \times 10^{-4}$ | $2 \times 10^{-4}$ | $10^{-4}$ |
| SHP-1 deactivation rate ($\beta$) | 0.503 | 1 | $6 \times 10^{-3}$ |

In our study, we aimed to understand how variations in parameter values impact the response of T cell activation within our model. The results of this analysis revealed distinct patterns of influence that various parameters exert on the model’s output, depending on the specific signaling conditions and feedback mechanisms considered.

### Key Findings

#### Phosphorylation Rate (*ϕ*) Influence

In KPR, KPR with Negative Feedback, KPR with Limited Signaling, and, KPR with Limited and Sustained Signaling, phosphorylation rate showed a positive PRCC value, indicating a significant positive correlation with T cell activation outcomes. This suggests that higher phosphorylation rates amplify TCR signaling, enhancing downstream immune responses such as cytokine production and T cell proliferation. Therefore, phosphorylation rate emerges as a critical parameter, with its modulation potentially influencing the strength and efficiency of T cell-mediated immunity. This finding underscores the importance of phosphorylation in regulating T cell activation dynamics.

#### Ligand/receptor binding Rate (*κ*) Impact

For the majority of models, ligand/receptor binding has a small, positive effect on T cell activation, indicating that while it contributes to activation, it is not the primary driver of the model’s output. This suggests that ligand binding serves as an initial trigger for activation, but other downstream factors are likely essential to achieve a robust or sustained response.

#### Off Rate (*v*) Impact

In case of KPR and occupancy, results indicate a small negative effect of the off-rate (*v*) on T cell activation. This is obvious as a higher *v* leads to a greater likelihood of the pMHC-TCR complex dissociating before reaching the fully modified, signaling-competent state *C*_*N*_, thereby diminishing the activation signal. However in KPR with limited and KPR with sustained signaling large negative values for Ω and *ξ* are found. This indicates that increasing Ω shortens the time TCRs remain in the signaling state, and *ξ* leads to faster deactivation of the signaling state. In other complex models the off rate (*v*) does not seem to impact the activation.

#### Phosphatase Efficiency (*γ*) in Feedback Scenarios

In the KPR model with negative feedback, the PRCC results indicate a competition between phosphatase efficiency and phosphorylation rate. The negative PRCC for phosphatase efficiency suggests that higher efficiency strengthens the feedback mechanism *S*, reducing activation, while the positive PRCC for phosphorylation rate shows that faster phosphorylation promotes activation. This balance underscores the model’s regulation of T cell signaling.

#### Amplification and Inhibition Parameters (*δ, σ* and *µ*)

In the KPR model with IFF and KPR with IFF and limited signaling, we found positive PRCC values for amplification parameters *σ* and *δ* in the enhancing pathway have positive values, indicating that increasing these parameters boosts T cell activation. Conversely, negative PRCC values for the inhibition parameter *µ* in the inhibitory pathway suggest that they suppress the signal. This balance between activation and inhibition allows fine-tuned control of the T cell response.

Also, positive PRCC for the total concentration of ligands in T cell models implies that an increase in ligand concentration enhances T cell activation. This suggests that higher ligand availability promotes the likelihood of TCRs reaching the signaling-competent state, thereby strengthening the activation response. Essentially, the model indicates that ligand concentration is a key driver of activation, with more ligands leading to a stronger or more sustained T cell response.

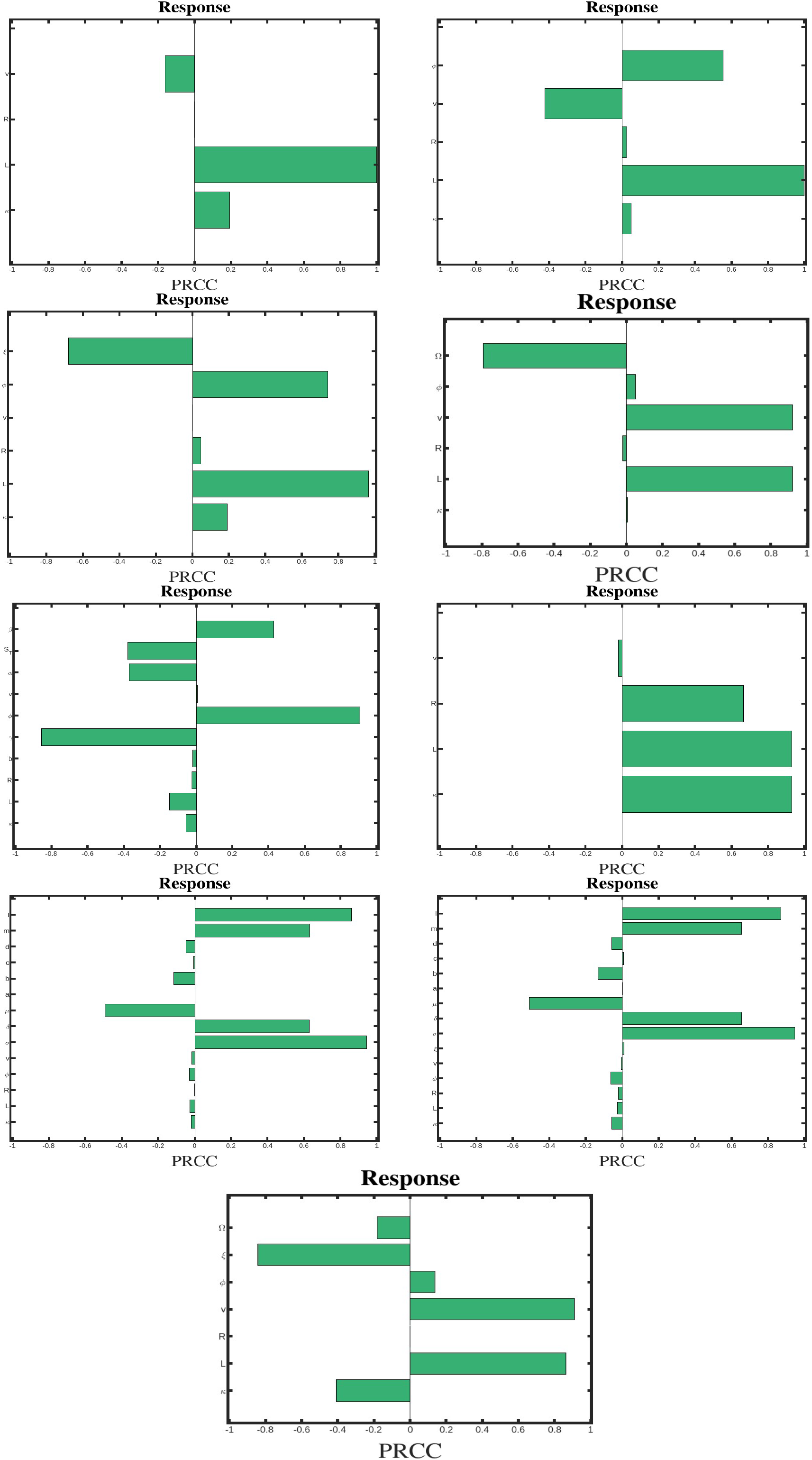

## 5 Parameter Dependency

The outcomes of our phenotypic models are highly dependent on the choice of parameters, with sensitivity analysis indicating that certain parameters have a more pronounced impact on the results. This underscores the importance of thoroughly exploring the process of parameter derivation.

Traditionally, 3D techniques have been employed to understand the interaction between TCR and pMHC. Techniques such as Surface Plasmon Resonance (SPR), Single Molecule Fluorescent Microscopy (SMFM), and Fluorescence Resonance Energy Transfer (FRET, in solution) are commonly used in these studies. These methods measure changes in molecular properties, such as refractive index or fluorescence, in response to TCR-pMHC binding events in solution. They provide crucial information on the thermodynamics, kinetics, and binding affinities of TCR-pMHC interactions, elucidating the molecular properties that govern binding events in bulk environments. The results produced by these 3D techniques have led to the formulation of several hypotheses and theoretical models to explain the observed outcomes. These models include kinetic proofreading [7, 53], TCR occupancy [54], serial triggering [12], optimal dwell-time [13], and rebinding [55].

One of the major limitations of these 3D techniques is their inability to accurately replicate the natural environment of T cell activation. In these methods, one of the interacting partners (e.g., the TCR or the pMHC) is immobilized on the sensor chip surface, while the other is introduced as a mobile analyte. However, in reality, the TCR is tethered to the T cell surface alongside CD3, and it engages with membrane-bound pMHC at the two-dimensional (2D) APC junction. Therefore to mimic the natural environment for T cell activation in recent years, various two-dimensional (2D) methods have been developed to directly assess the kinetics of TCR-pMHC interactions on live T cells.

In 2D models, TCR and pMHC molecules interact on a flat surface, closely replicating the physiological conditions where TCRs are rooted to the T cell surface and engage with pMHCs presented by APCs. Examples of 2D techniques include supported planar lipid bilayers, micropipette adhesion frequency assays, thermal fluctuation assays, and single-molecule microscopy. These methods enable the direct observation of TCR-pMHC interactions on cell surfaces or artificial lipid bilayers, providing a more accurate representation of the natural cellular environment.

The primary experimental findings from these 2D and 3D techniques include the values for *k*_off_, *k*_on_, and the *k*_*D*_ of the TCR-pMHC interaction. First of all, in 2D methods, affinity is measured in *µm*^4^, with *k*_on_ (*µm*^4^*s*^−1^) and *k*_off_ (s^−1^), while in 3D methods, affinity is measured in M (*K*_*D*_), with *k*_on_ (M^−1^s^−1^) and *k*_off_ (s^−1^). Comparing *k*_on_, *k*_off_, and *k*_*D*_ values obtained from 2D and 3D methods reveals notable differences, including a broader range of 2D affinities and on-rates, along with significantly faster 2D off-rates for various pMHCs with different functions. The key distinctions between 2D and 3D parameter values include:

1. The interactions between antigenic peptide-major histocompatibility complexes (pMHC) and TCRs exhibited both high affinity and rapid association-dissociation kinetics.
2. Ligands of different potencies displayed a broad spectrum of affinities and binding rates, highlighting their diverse dynamic ranges.

The variations observed in parameter values between 2D and 3D settings can be attributed to a combination of physical and biological factors. The unique physical chemistry of the binding site is usually clarified using crystal structures and intrinsic interaction parameters obtained from purified recombinant molecules. However, the spatial constraints of different environments can influence binding dynamics differently. From a biological perspective, the differences are greatly influenced by molecular interactions occurring within the natural membrane environment. The parameter values we used were sourced from four distinct references that employed 3D techniques. It may be difficult to directly apply parameters from 2D techniques due to unit differences, as in case of 2D techniques affinity is typically measured in *µm*^4^ and *k*_on_ in *µm*^4^ · *s*^−1^, while parameters like association and phosphorylation rates are usually measured as *k*_on_ = L/mol · s^−1^ or *k*_on_ = concentration^−1^ · time^−1^. Hence, the sources from which we have taken our parameters are based on 3D models, ensuring consistency in unit measurement.

## 6 Discussion

T cells use their TCRs to detect peptides displayed by pMHC molecules, distinguishing between low-affinity self-peptides and high-affinity foreign peptides. Only strongly binding foreign peptides trigger a full immune response. This study evaluates nine models of T cell activation to replicate key immune response features like specificity and antigen discrimination.

We examined the long-term behavior of various systems, focusing on steady states and convergence properties. We found that most systems, including the Occupancy Model, KPR, and KPR variants, exhibit convergence to a unique steady state, confirmed using the Deficiency Zero Theorem. However, the KPR model with negative feedback does not converge to a unique steady state, consistent with prior findings [25].

The Occupancy Model links T-cell activation to the number of TCR-pMHC complexes but fails to capture antigen discrimination due to a constant maximum response *E*_max_. The Kinetic Proofreading (KPR) model addresses this by incorporating time dependency, allowing only ligands with sufficient binding duration to activate T cells, explaining antigen discrimination. Variants of KPR further enhance immune response accuracy: limited signaling introduces temporal restrictions for optimal responses, sustained signaling reflects extended signaling post-detachment, and negative feedback prevents excessive activation with bell-shaped dose-response curves. Models with incoherent feedforward (IFF) loops optimize sensitivity and specificity, particularly the KPR with IFF and limited signaling, which balances high-affinity interactions with limited low-affinity responses.

Our mathematical and numerical analysis of these models uncovered several novel insights. In the sustained signaling model, we identified an optimal response across all dissociation times concerning ligand concentration. For the negative feedback signaling model, an optimal response emerged with respect to dissociation time, aligning with the predictions of [25], though the optimum was not strongly pronounced. In the IFF model, we observed intersections in dose-response curves at different ligand concentrations. Notably, ligands with longer dissociation times elicited the strongest responses before the peak, contrasting with the findings of [31]. Furthermore, in the IFF model with limited signaling, we discovered a previously unreported bimodal response, adding a new dimension to the existing understanding of ligand-induced signaling dynamics.

Our sensitivity analysis, using LHS-PRCC methods, revealed that the phosphorylation rate (*ϕ*) significantly influences T cell activation dynamics. Phosphorylation rate strongly enhances activation, while higher off-rates reduce signaling duration by increasing TCR-pMHC dissociation. Phosphatase efficiency (*γ*) modulates feedback regulation, balancing activation and suppression. Amplification and inhibition parameters (*δ, λ, µ*) fine-tune the immune response, and ligand concentration emerges as a key driver of activation. These findings highlight the importance of precise parameter selection for accurate modeling.

Given that the sensitivity analysis revealed that model outcomes are highly dependent on parameter choice, we compared parameter values from traditional 3D techniques to those from emerging 2D methods, which better replicate physiological conditions. The comparison showed that 2D techniques provide a broader range of affinities and faster off-rates, reflecting the dynamic and spatially constrained nature of the T-cell environment.

These findings underscore the critical importance of employing context-appropriate experimental methodologies to enhance the accuracy and relevance of parameter derivation in phenotypic modeling. Since the outcomes of our models are inherently dependent on parameter values, achieving precise and biologically realistic parameter estimates is essential for generating reliable and robust predictions. A systematic and rigorous approach to parameter calibration will significantly improve the validity and applicability of model outcomes.

In conclusion, this study establishes the efficacy of KPR-based models in simulating T-cell activation, offering a comprehensive evaluation of their strengths while identifying key areas for refinement. Specifically, our findings emphasize the need for improved experimental parameterization to enhance model precision. By addressing these limitations, this research provides a foundation for refining T-cell activation models to better capture the complexity of immune response dynamics. These insights contribute not only to the field of computational immunology but also to the broader understanding of T-cell-mediated immunity, paving the way for more accurate predictive tools and novel therapeutic strategies.

## A Appendix A

**Fig. 9.**
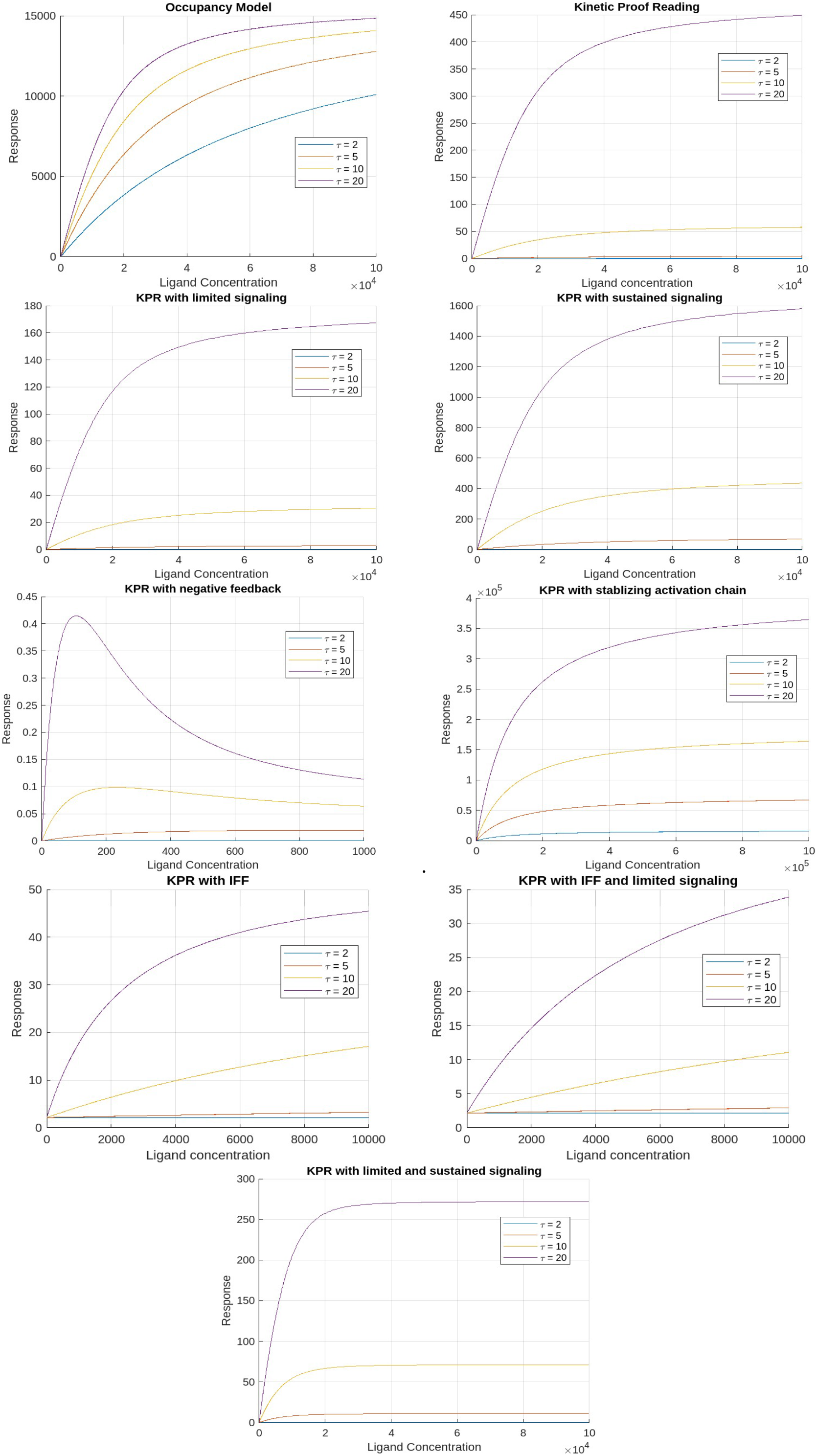
Dose-response predictions for ligands of increasing dissociation time ((*τ*) (for minimum values of parameters)

**Fig. 10.**
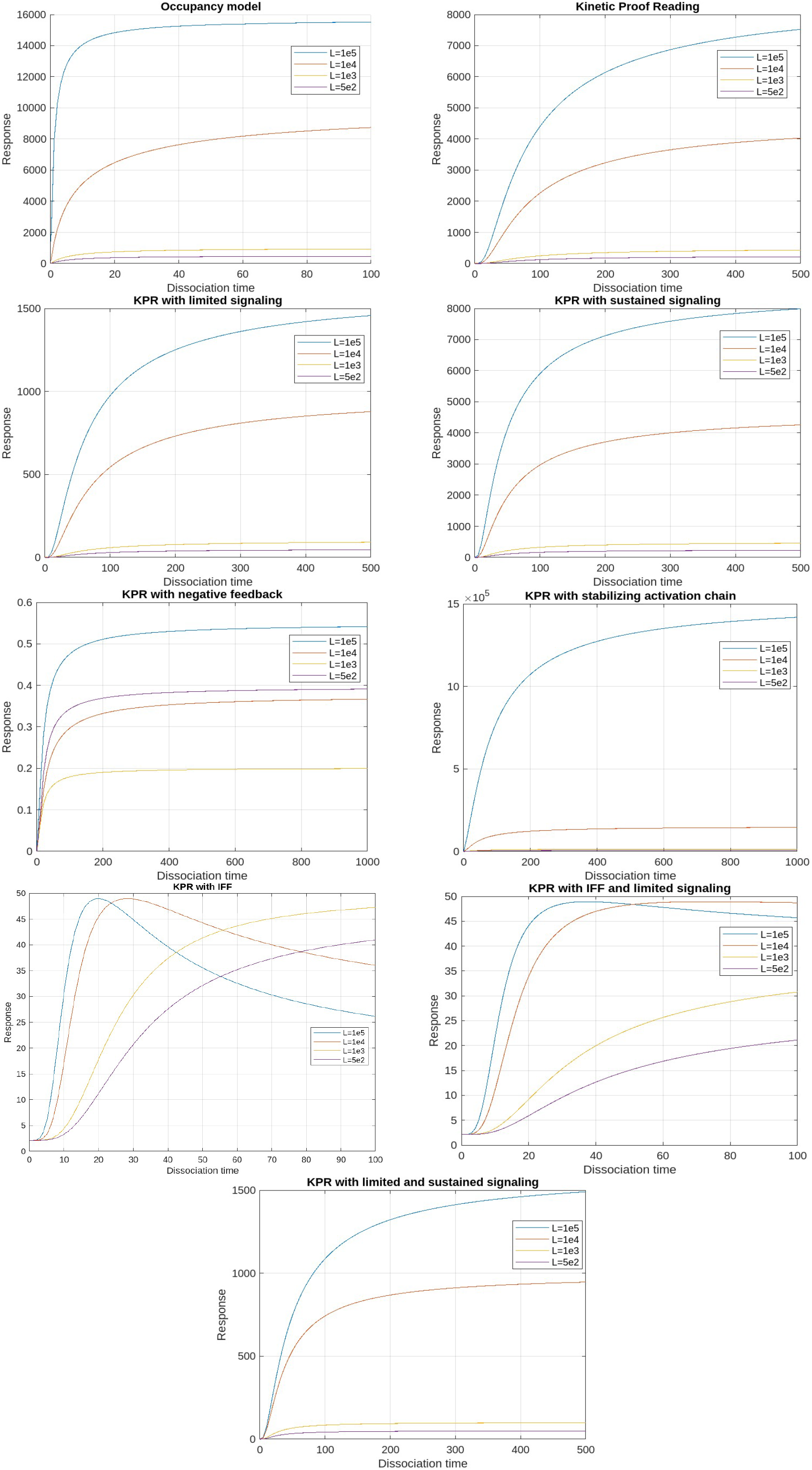
T cell activation as a function of ligand dissociation time (*τ*) at a fixed ligand dose (for minimum values of parameters).

## B Appendix B

In this study, we also investigated the late-time behavior of our models, focusing on the existence and uniqueness of steady states and convergence properties as *t* → ∞. Specifically, we address the question: Does each choice of parameters yield a unique steady state, and do all solutions converge to this steady state in the long term? Our findings are as follows:

1. For system (1) (Occupancy Model), the answer is affirmative. Given that this system is effectively one-dimensional, the proof of convergence to a unique steady state is straightforward.
2. For systems (2) and (6) (KPR and KPR with stabilizing activation chain), convergence to a unique steady state is also established, applying the theorem by Sontag, as referenced in prior communications.
3. For systems (3), (4), and (9) (KPR with limited signaling, sustained signaling, and both), convergence to a unique steady state is confirmed based on the application of the Deficiency Zero Theorem, as demonstrated in the following sections.
4. For the system (5) (KPR with negative feedback), however, the answer is negative, which aligns with the earlier findings.
5. Lastly, for systems (7) and (8) (KPR with IFF, and limited signalling), the answer is affirmative. The convergence to a unique steady state is proven below.

### B.1 Kinetic Proofreading

Within this framework, a pMHC ligand (L) can bind reversibly to a TCR receptor (R), resulting in the formation of a pMHC-TCR complex denoted as *C*_0_. Once formed, this TCR-pMHC complex (*C*_0_) undergoes a series of biochemical changes to achieve a signaling-competent state represented as *C*_*N*_. If a pMHC detaches from a TCR at any intermediate stage, all modifications are instantly undone, causing the TCR to return to its original, unmodified state. T cell activation is directly proportional to the quantity of TCRs in the *C*_*N*_ state.

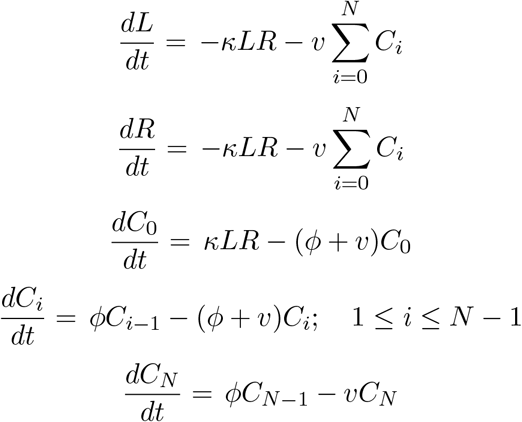

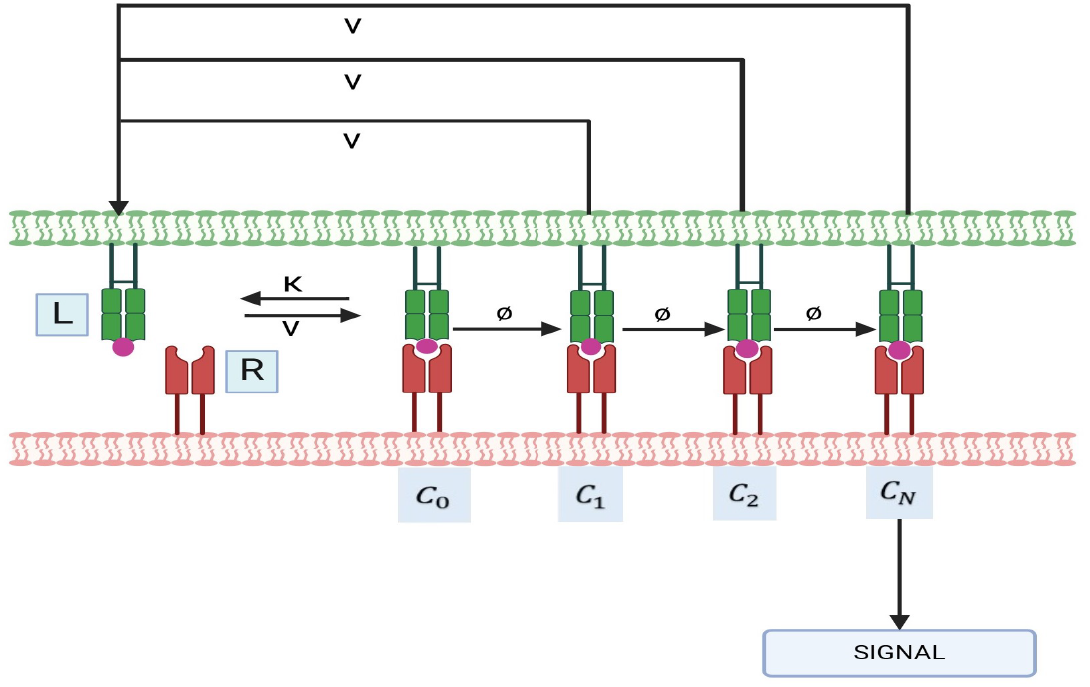

#### Mathematical Formulation

The above equations can be rewritten as:

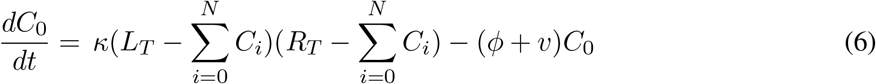

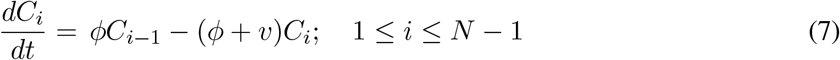

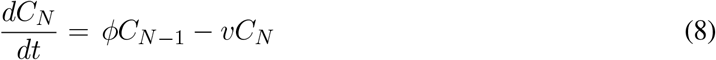

Here *R*_*T*_ and *L*_*T*_ are the total concentrations of the receptor and the ligand.

From equations (6)-(8), it can be deduced that if 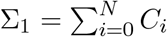

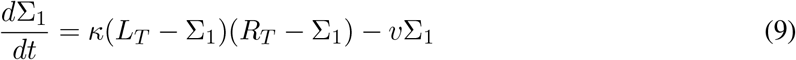

##### Lemma B.1.

*Let* (*C*_0_(*t*), *C*_1_(*t*), …, *C*_*N*_ (*t*)) *be a solution of (6)-(8) contained in the closure K of the biologically relevant region. Then any ω-limit point* 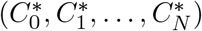 *of this solution is contained in the interior of K. In particular, any steady state is contained in the interior of K*.

*Proof*. For the proof we use Lemma 2.1. It follows from equation (9) that 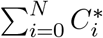 is strictly less than*L*_*T*_ and *R*_*T*_. It then follows from (6) that 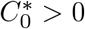. This in turn implies using (7) and (8) that 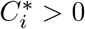 for 1 ≤ *i* ≤ *N*.

#### Stability of the solutions

In case of KPR, the binding reaction is of the form *L* + *R* = *C*_0_. The bound receptor *C*_0_ undergoes a series of phosphorylations to reach the signalling competent state *C*_*N*_. Each *C*_*i*_ can decay releasing *L, R* and the phosphate groups. The network is weakly reversible. In our system, there are *n* = *N* + 2 complexes, and it contains only a single linkage class, which means *l* = 1. To show that the deficiency of the above weakly reversible network is zero it will be sufficient to show that the rank of the above system is *s* = *N* + 1.

There are *N* + 2 complexes {*L* + *R, C*_0_, *C*_1_, *C*_2_, …., *C*_*N*_}.

The stoichiometric matrix is given by:

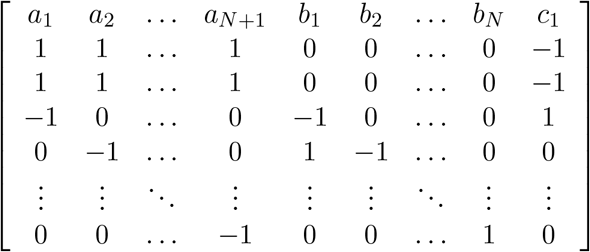

The first *N* + 1 columns of this matrix are linearly independent. Hence for this model *s* ≥ *N* + 1. Hence *δ* ≤ (*N* + 2) − (*N* + 1) − 1 = 0. Since *δ* is always non-negative this implies that *δ* = 0.

### B.2 Kinetic proofreading with limited signaling

This model extends the kinetic proofreading concept, suggesting that once a TCR reaches the signaling-competent state *C*_*N*_, the bound TCR shifts to a non-signalling state *C*_*N*+1_ at a rate *ξ*.

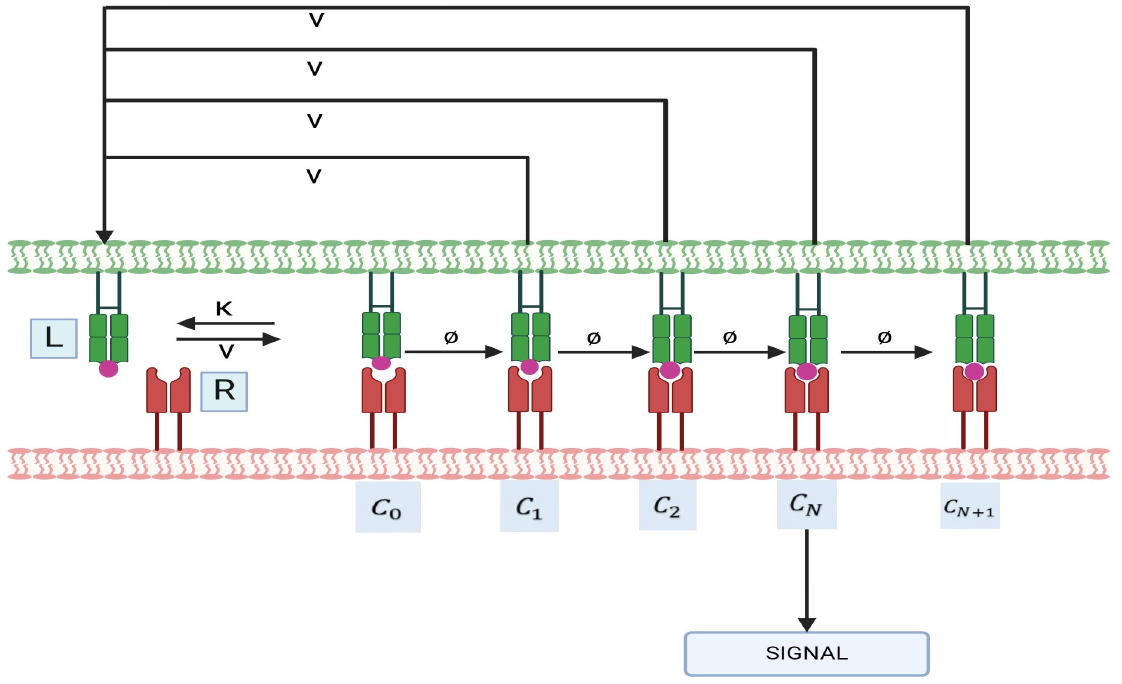

#### Mathematical formulation

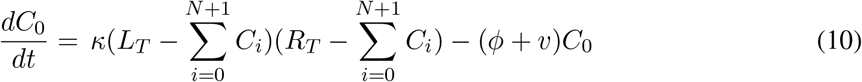

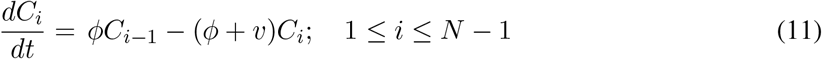

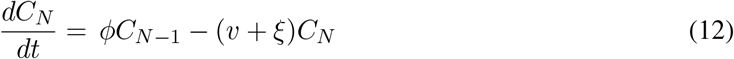

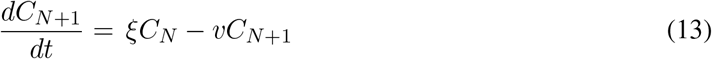

Here *R*_*T*_ and *L*_*T*_ are the total concentrations of receptor and the ligand. Let 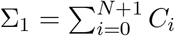. Then, from Equations (10)-(13), we have:

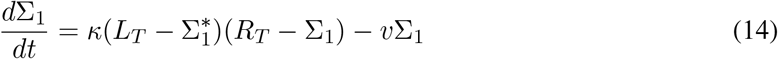

##### Lemma B.2.

*Let* (*C*_0_(*t*), *C*_1_(*t*), …, *C*_*N*+1_(*t*)) *be a solution of (10)-(13) contained in the closure K of the biologically relevant region. Then an ω-limit point* 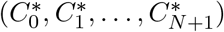 *of this solution is contained in the interior of K. In particular, any steady state is contained in the interior of K*.

*Proof*. For the proof we use Lemma 2.1. It follows from equation (14) that 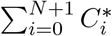 is strictly less than *L*_*T*_ and *R*_*T*_. It then follows from (10) that 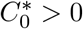. This in turn implies inductively, using (10)-(13) that 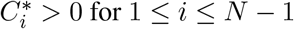.

We have proved the deficiency of this weakly reversible network is zero as in section 3.

### B.3 Kinetic proofreading with sustained signalling

This model extends the KPR scheme by integrating experimental findings that indicate signalling-competent TCRs can maintain signaling for a defined time frame, even once pMHC unbinding occurs. In this framework, T-cell receptors (TCRs) in the signalling-capable state *C*_*N*_ persist in signaling for a designated period (*T*) following the detachment of pMHC, subsequently returning to their baseline state at a rate *λ*.

#### Mathematical Formulation

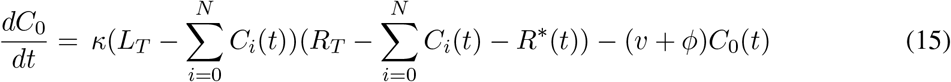

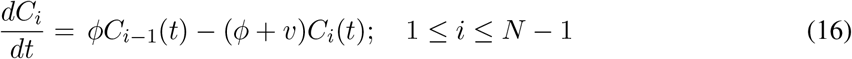

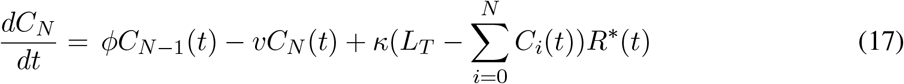

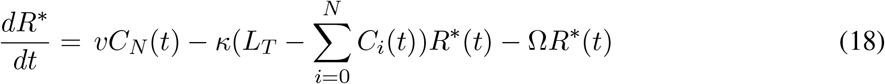

Here *R*_*T*_ and *L*_*T*_ are the total concentrations of receptors and the ligand.

If 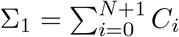 then it follows from equations (15)-(18) that:

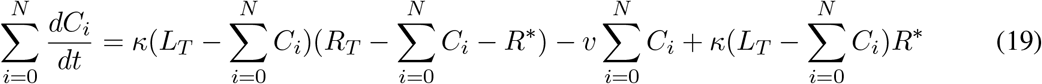

and

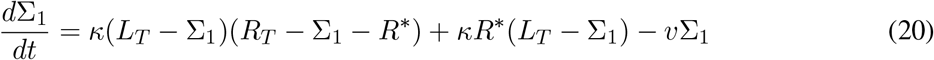

##### Lemma B.3.

*Let ((R*^∗^(*t*), *C*_0_(*t*), *C*_1_(*t*), …, *C*_*N*_ (*t*)) *be a solution of (15)-(18) contained in the closure K of the biologically relevant region. Then any ω-limit point* 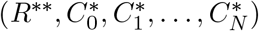 *of this solution is contained in the interior of K. In particular, any steady state is contained in the interior of K*.

*Proof*. For the proof we use Lemma 2.1. It follows from (20) that 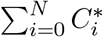 is strictly less than *L*_*T*_. Suppose now that 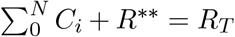. Then it follows from (15) that 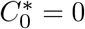. This implies, using (16) that 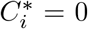 for all 1 ≤ *i* ≤ *N* − 1. The sum of (17) and (18) then implies that *R*^∗∗^ = 0. Substituting this back in (17) shows that 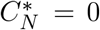. Putting these facts together we see that 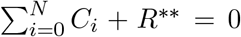, contradicting our assumption. Thus in fact Σ_1_ + *R*^∗∗^ *< R*_*T*_. Once this has been established it follows from (15) that 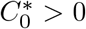. Then (16) implies that 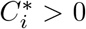 for 1 ≤ *i* ≤ *N* − 1. From (17) we can conclude that 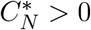 and from (18) that *R*^∗∗^ *>* 0.

#### Stability of the Solutions of system

In the case of kinetic proofreading (KPR) with sustained signaling, the binding reaction follows *L* + *R* = *C*_0_, where the bound receptor complex *C*_0_ undergoes a sequence of phosphorylation steps to reach the signaling-competent state *C*_*N*_. Unlike in limited signaling, here the TCRs continue to signal even after pMHC dissociates. The rate at which these unbound yet signaling-competent TCRs revert to their unmodified state is governed by a factor Ω. In this model, T cell activation depends on both *C*_*N*_ and *R*^∗^. Each intermediate state *C*_*i*_ can decay, releasing *L, R, R*^∗^, and phosphate groups, making the network weakly reversible.

Our system consists of *n* = *N* + 3 complexes with a single linkage class (*l* = 1). To establish that the deficiency of this weakly reversible network is zero, it suffices to show that its rank is *s* = *N* + 2.

The network comprises *N* + 3 complexes: {*L* + *R, C*_0_, *C*_1_, *C*_2_, …, *C*_*N*_, *L* + *R*^∗^}.

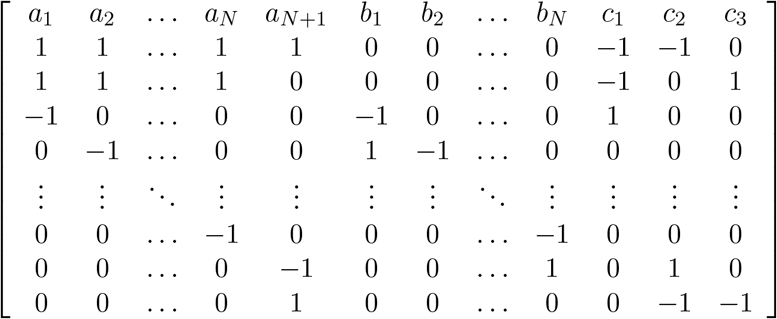

The first *N* + 2 columns of this matrix are linearly independent. Hence for this model *s* ≥ *N* + 2. Hence *δ* ≤ (*N* + 3) − (*N* + 2) − 1 = 0. Since *δ* is always non-negative this implies that *δ* = 0.

### B.4 Kinetic Proofreading with Stabilizing Activation Chain

The model indicates that KPR complexes enhance the stability of foreign peptides while reducing their affinity for self-peptides. This selective strengthening and weakening of the *C*_*i*_ complexes, as well as differences in activation timing, are represented through changes in the respective rate constants *v*(*i*); (*i* = 0, 1, …, *N*) and *ϕ*(*i*); (*i* = 0, 1, …, *N* − 1) as the proofreading process advances.

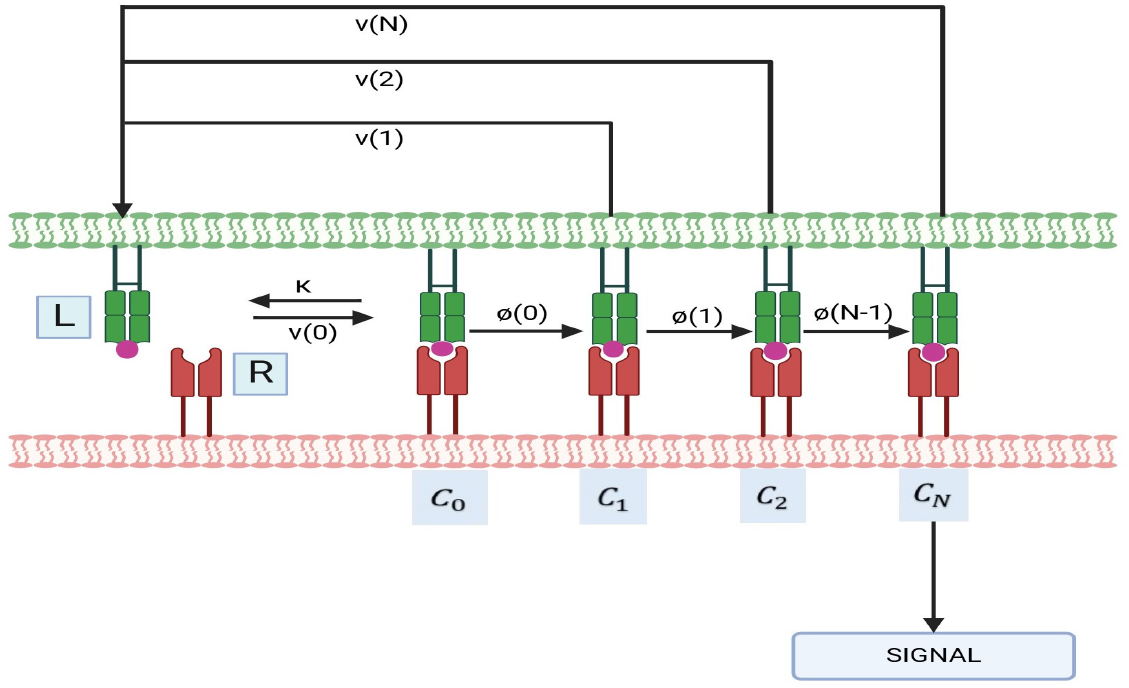

#### Mathematical formulation

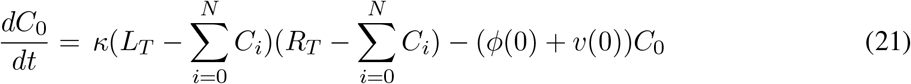

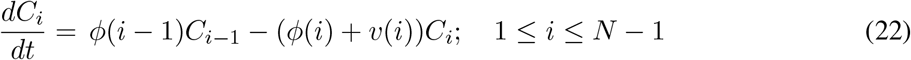

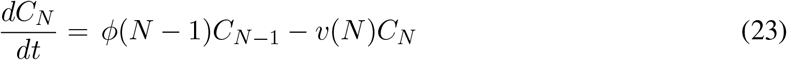

In this model for simplicity *v*(0) is denoted as *v*(= 1*/τ*) which is dissociation time, and *ϕ*(0) denotes the propagation rate for the first step *C*_0_ → *C*_1_. Now for next steps these rates are given by

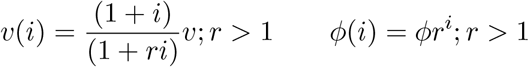

Let 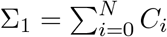. From Equations (16)-(18), we get:

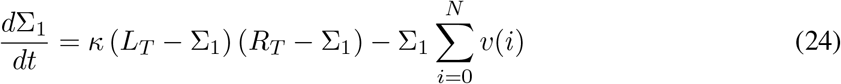

##### Lemma B.4.

*Let* (*C*_0_(*t*), *C*_1_(*t*), …, *C*_*N*_ (*t*)) *be a solution of (21)-(23) contained in the closure K of the biologically relevant region. Then any ω-limit point* 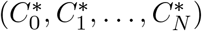 *of this solution is contained in the interior of K. In particular, any steady state is contained in the interior of K*.

*Proof*. For the proof we use Lemma 2.1. It follows from (24) that 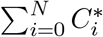 is strictly less than *L*_*T*_ and *R*_*T*_. It then follows from (21) that 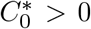. This in turn implies using (22) and (23) that 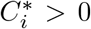 for 1 ≤ *i* ≤ *N*.

#### Stability of the solutions

In the case of KPR with a stabilizing activation chain, the binding reaction follows *L*+*R* = *C*_0_, where the bound receptor complex *C*_0_ undergoes a sequence of phosphorylation steps to reach the signaling-competent state *C*_*N*_. T cell activation is determined by *C*_*N*_. Each intermediate state *C*_*i*_ can decay, leading to the release of *L, R*, and phosphate groups, making the network weakly reversible.

Our system consists of *n* = *N* + 2 complexes with a single linkage class (*l* = 1). To establish that the deficiency of this weakly reversible network is zero, it suffices to show that its rank is *s* = *N* + 1.

The network comprises *N* + 2 complexes: {*L* + *R, C*_0_, *C*_1_, *C*_2_, …, *C*_*N*_}.

The stoichiometric matrix is given by:

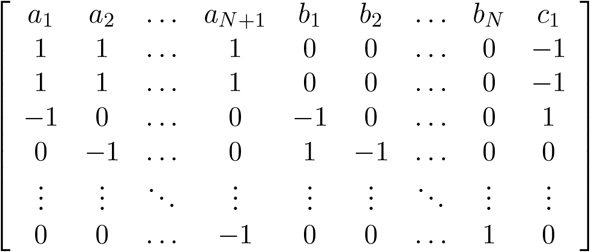

The first *N* + 1 columns of this matrix are linearly independent. Hence for this model *s* ≥ *N* + 1. Hence *δ* ≤ (*N* + 2) − (*N* + 1) − 1 = 0. Since *δ* is always non-negative this implies that *δ* = 0.

### B.5 Kinetic Proofreading with Limited and Sustained Signaling

This model is a combination of two models KPR with limited and KPR with sustained signaling.

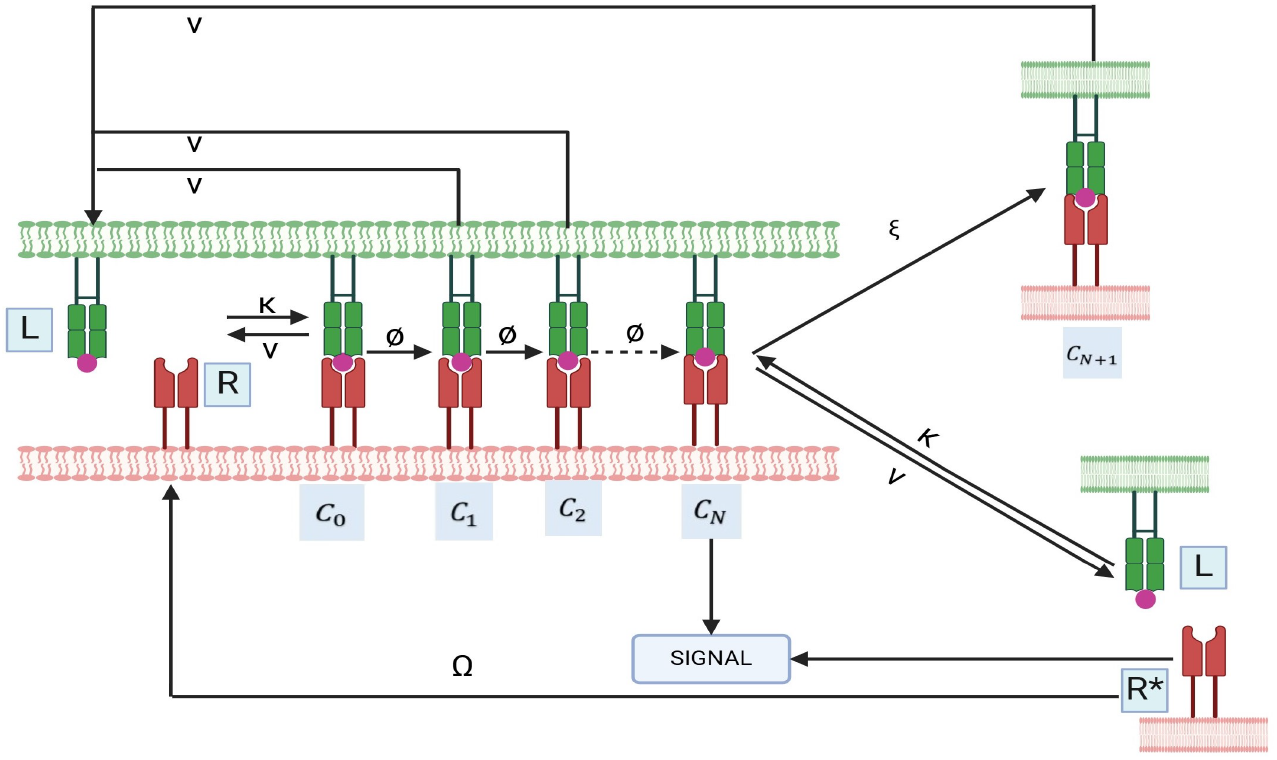

#### Mathematical formulation

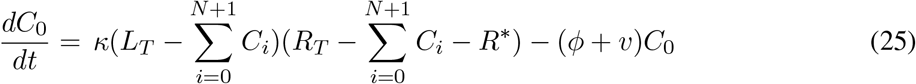

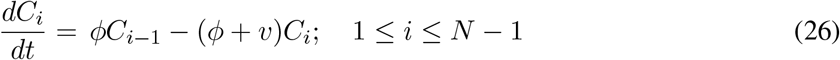

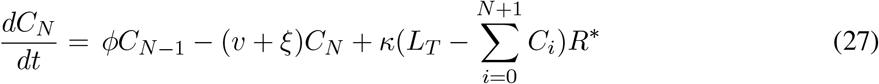

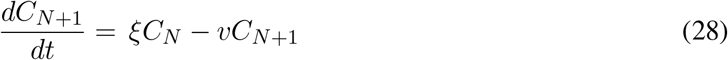

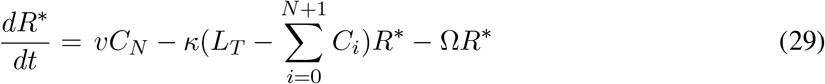

Define 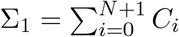. From equations (25)-(28) it follows that:

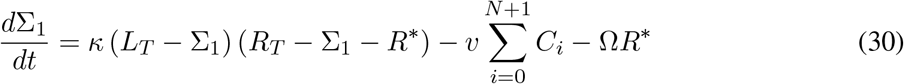

##### Lemma B.5.

*Let ((C*_0_(*t*), *C*_1_(*t*), …, *C*_*N*+1_(*t*), *R*^∗^(*t*)) *be a solution of (25)-(28) contained in the closure K of the biologically relevant region. Then any ω-limit point* 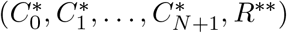 *of this solution is contained in the interior of K. In particular, any steady state is contained in the interior of K*.

*Proof*. For the proof we use Lemma 2.1. It follows from (30) that 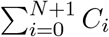 is strictly less than *L*_*T*_. Suppose now that 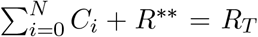. Then it follows from (25) that 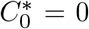. This implies, using (26) that 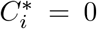 for all 1 ≤ *i* ≤ *N* − 1. The sum of (27), (28) and (29) implies that *R*^∗∗^ = 0. Substituting this back into (28) gives *C*_*N*+1_ = 0 and substituting this into (17) gives *C*_*N*_ = 0. Putting these facts together we see that 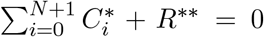, contradicting our assumption. Thus in fact 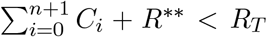. Once this has been established it follows from (25) that 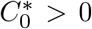. Then (26) implies that 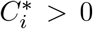 for 1 ≤ *i* ≤ *N* − 1. From (27) we can conclude that 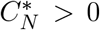, from (28) that 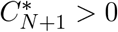 and from (29) that *R*^∗∗^ *>* 0.

#### Stability of the solutions

In case of KPR with limited and sustained signalling the binding reaction is of the form *L* + *R* = *C*_0_. The bound receptor *C*_0_ undergoes a series of phosphorylations to reach signalling competent state *C*_*N*_. In this model, T cell activation is determined by both *C*_*N*_ and *R*^∗^. Each *C*_*i*_ can decay releasing *L*,*R, R*^∗^, and the phosphate groups. The network is weakly reversible. In our system we have *n* = *N* + 4 complexes; there is only one linkage class i.e., *l* = 1. To show that the deficiency of the above weakly reversible network is zero. It will be sufficient to show that the rank of the above system is *s* = *N* + 3.

There are *N* + 4 complexes {*L* + *R, C*_0_, *C*_1_, *C*_2_, …., *C*_*N*+1_, *L* + *R*^∗^}.

The stoichiometric matrix is given by:

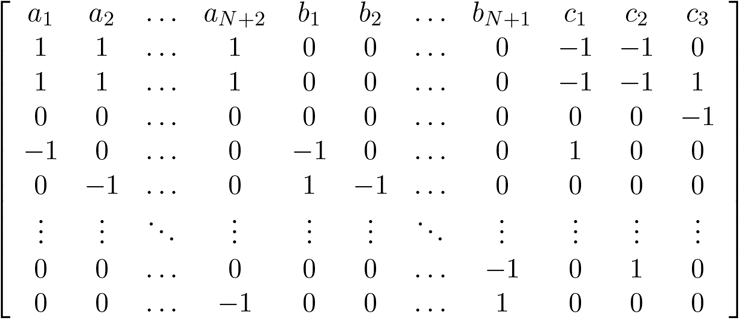

The first *N* + 3 columns of this matrix are linearly independent. Hence for this model *s* ≥ *N* + 3. Hence *δ* ≤ (*N* + 4) − (*N* + 3) − 1 = 0. Since *δ* is always non-negative this implies that *δ* = 0.

### B.6 Kinetic proofreading with limited signaling and incoherent feed forward loop

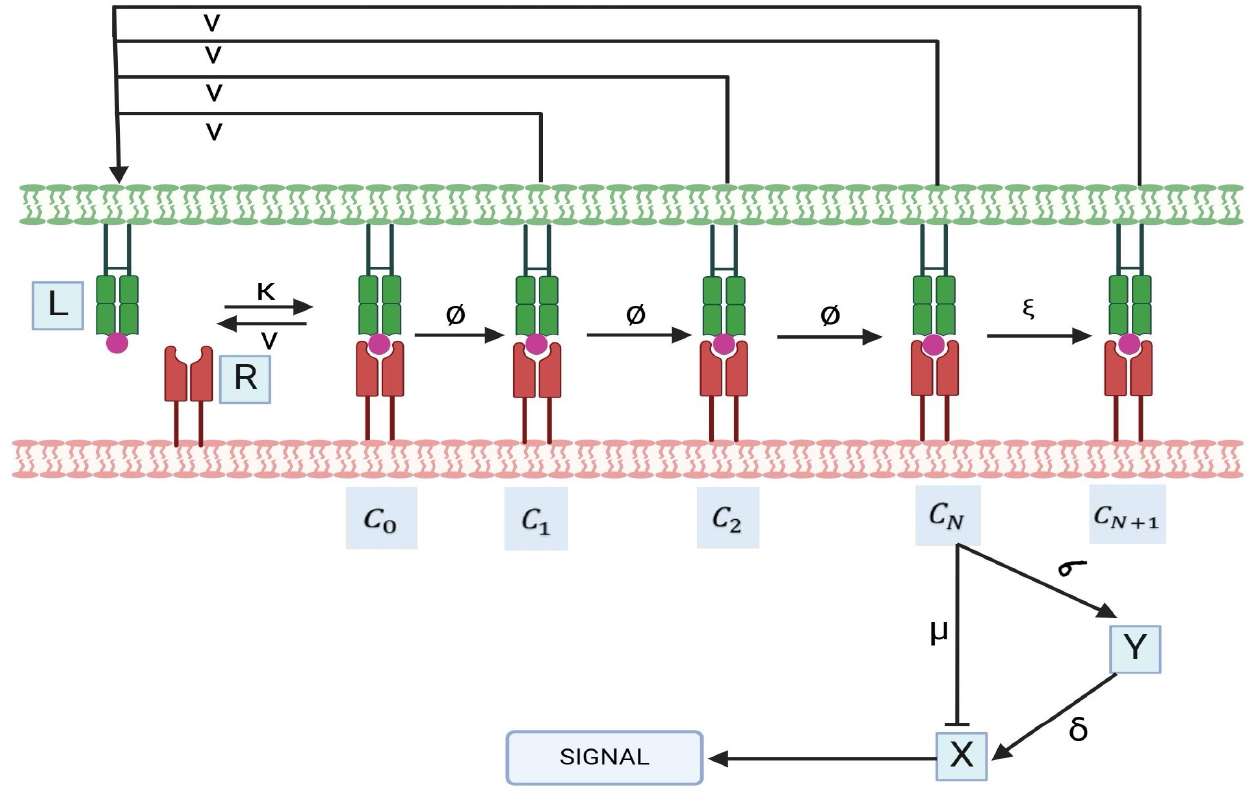

This model is an extension of KPR with an incoherent feed forward loop, it has been assumed that signalling is limited and after reaching the signalling competent state the bound TCR transits to a non-signalling transit state.

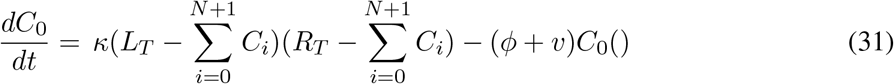

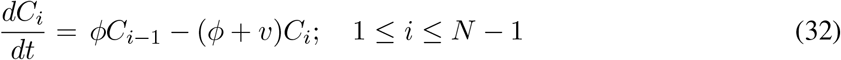

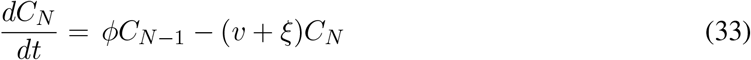

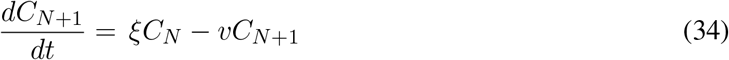

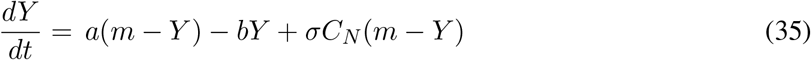

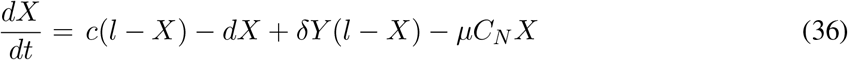

We aim to demonstrate that the system, an extension of the kinetic proofreading (KPR) model, is globally asymptotically stable. The system consists of two sets of variables: *Z*_1_, consisting of the *C*_*i*_, which satisfies a closed system of equations

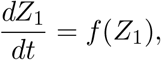

and *Z*_2_, consisting of *X* and *Y*, governed by

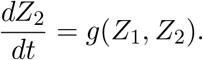

It has already been proven that the system for *Z*_1_ has a unique steady state, and that all solutions for *Z*_1_ converge to this steady state as *t* → ∞. In other words, as *t* → ∞ each *C*_*i*_(*t*) converges to some 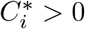. Consider any *ω*-limit point of a solution (*C*_*i*_, *X, Y*) of (31)-(36) and the solution starting at that point for *t* = 0. It lies entirely in the *ω*-limit set of that solution and so *C*_*i*_ has the constant value 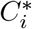 for all *i*. This means that this solution satisfies the equations obtained from (35) and (36) by replacing *C*_*N*_ by 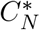. Equation (35) is an equation for *Y* alone and so it it easy to see that the solution converges to 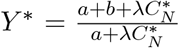 as *t* → ∞. Now we can again pass to a solution starting at an *ω*-limit point to see that for the resulting solution *X* solves the equation obtained from (36) by replacing *C*_*N*_ by 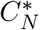 and *Y* by *Y*^∗^. Thus the solution converges to 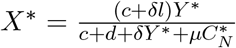 as *t* → ∞. Since these statements hold for all *ω*-limit points it follows that any solution of (31)-(36) converges to 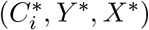 as *t* → ∞. For this system the unique positive steady state is globally asymptotically stable.

### B.7 Kinetic proofreading with incoherent feed forward loop

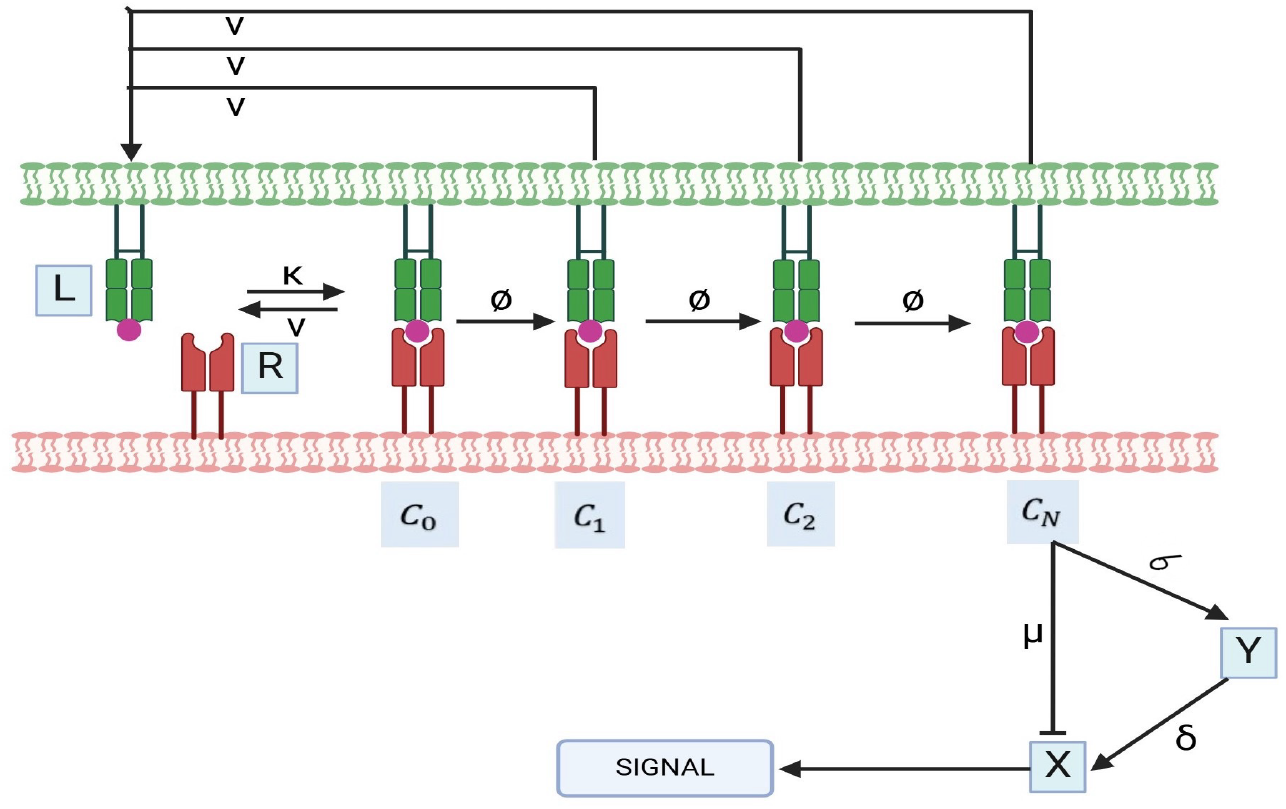

For this model there is a unique steady state in each stoichiometric compatibility class which is globally asymptotically stable. The proof of this is based on the fact that we have already proved global asymptotic stability for the kinetic proofreading model and otherwise the proof is just as in the previous example.

### B.8 Analytical observations on results

#### Occupancy model

Response as a Monotonic Function of Dissociation Time:

In the context of the occupancy model, we analyze the response by examining T cell activation as a function of receptor-ligand complex concentration, *C*, which is derived from the equilibrium concentrations of ligand and receptor. It satisfies the equation:

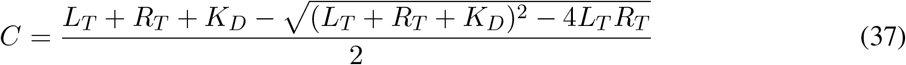

and is this model *C* is a measure of the T cell activation.

*C* is a solution to a quadratic equation of the form:

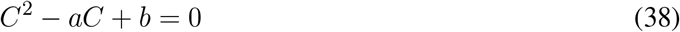

where

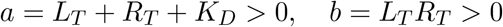

Only the negative square root gives a relevant solution since *C < L*_*T*_ and *C < R*_*T*_ and hence *C < a*. The above equation can be reformulated as:

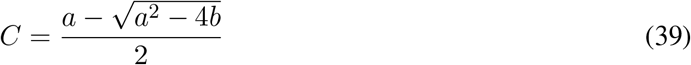

Analysis of the Solution

Differentiating the expression for *C* with respect to *K*_*D*_ gives

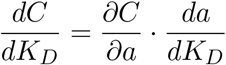

*C* is a decreasing function of *a* since

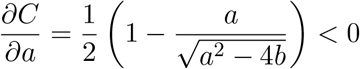

and combining this with

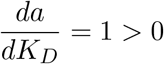

we have:

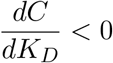

Relationship with Dissociation Time

The dissociation constant *K*_*D*_ is related to the dissociation rate constant *k*_off_ and the association rate constant *k*_on_:

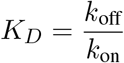

Additionally, the dissociation time *τ* is inversely proportional to the dissociation rate constant:

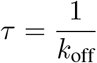

As the dissociation time *τ* increases, the dissociation rate constant *k*_off_ decreases, resulting in a decrease in *K*_*D*_. Given that *C* is a decreasing function of *K*_*D*_, it follows that *C* becomes an increasing function of the dissociation time *τ*.

This relationship demonstrates that as ligand-receptor binding stabilizes, resulting in a longer dissociation time, T cell activation increases, highlighting the importance of bond longevity in cellular activation responses.

#### Response as monotonic function of ligand concentration

To determine how *C* depends on *L*_*T*_, we differentiate *C* with respect to *L*_*T*_ :

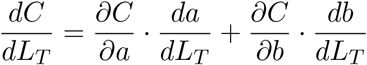

We have 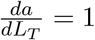 and 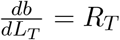. Differentiating *C* with respect to *b* gives:

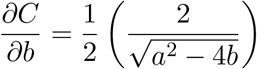

Sign of 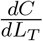

Analyzing the derivative 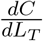:

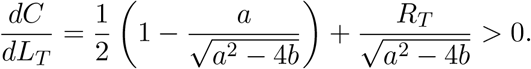

Thus, the equilibrium concentration of the pMHC-TCR complex (*C*) is an increasing function of the ligand concentration (*L*_*T*_). As the ligand concentration increases, the number of available binding sites increases, which directly enhances the formation of the complex (*C*). Therefore, in the occupancy model, the response function is positively dependent on the ligand concentration.

#### Response of KPR model as increasing function of dissociation time

In the kinetic proofreading model the response function is given by:

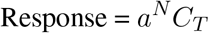

where

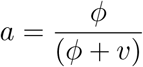

and

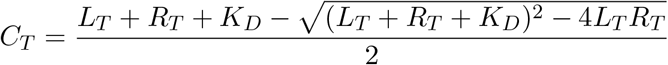

Similar to the argument in occupancy model it is clear that for this model *C*_*T*_ satisfies the same equation as *C* does in the occupancy model. Thus it is a decreasing function of *k*_off_, which follows that it is an is an increasing function of the dissociation time (*τ*).

#### Response of KPR model as increasing function of ligand concentration

That the response with respect to ligand concentration is monotone in the kinetic proofreading model can be attributed again to the fact that in this case *C*_*T*_ is increasing (similar to the argument in the case of the occupancy model) and *α* is constant. Thus *R* is increasing.

## C Mathematical Formulation and analysis

### C.1 Occupancy Model

In this model a pMHC ligand (L) can reversibly bind a T cell receptor (R) to form a pMHC-TCR complex (C).

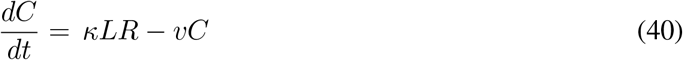

where

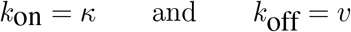

and at equilibrium 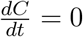 and hence

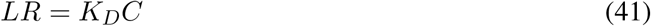

where 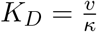

Also the total amount of ligand *L*_*T*_ and total amount of receptor TCR *R*_*T*_ are conserved quantities

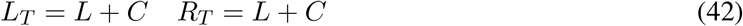

By inserting these in equation in (41) we get

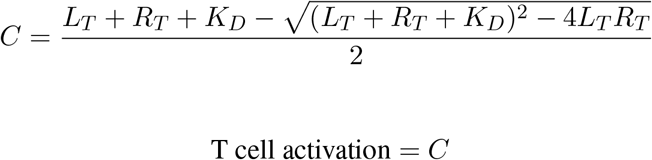

#### Calculating the *E*_max_

The *E*_max_ can then be found by finding *C* in the limit of *L*_*T*_ tending to infinity. In this limit, we have

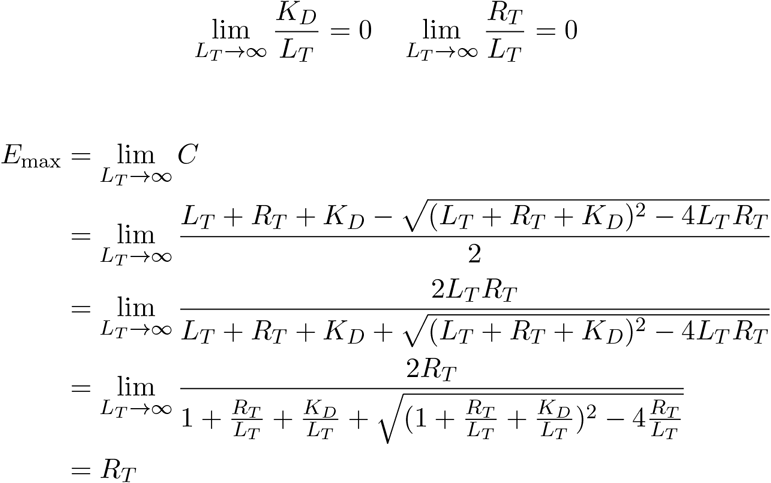

Hence

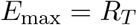

#### Calculation for *EC*_50_

At half the maximal response, we have *L*_*T*_ = *EC*_50_.

Also, T cell activation =

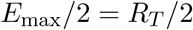

which implies

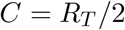

Using equation (41) we have

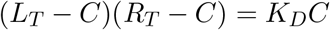

hence

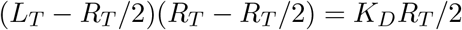

Rearranging we get

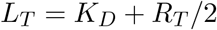

Hence

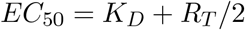

### C.2 Kinetic Proofreading

In this model, a pMHC ligand (L) can reversibly associate with a TCR receptor (R) to form a pMHC-TCR complex, denoted as *C*_0_. Once formed, this complex undergoes a series of biochemical modifications, progressively transitioning towards a signaling-competent state, labeled as *C*_*N*_. If the pMHC dissociates from the TCR at any intermediate stage, all modifications are rapidly undone, causing the TCR to revert to its original, unmodified state. The level of T cell activation is directly linked to the quantity of TCRs in the fully modified *C*_*N*_ state.

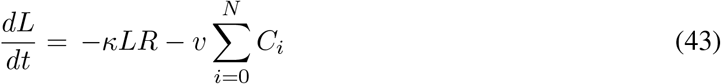

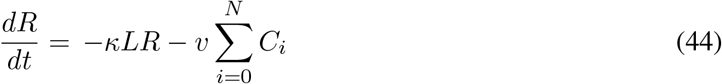

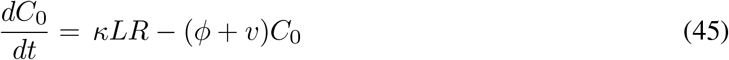

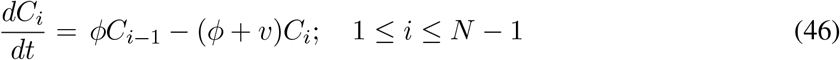

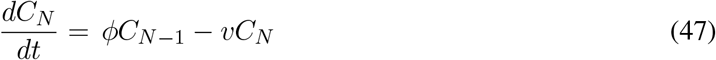

We have the total amount of ligand *L*_*T*_ and total amount of receptor TCR *R*_*T*_ as conserved quantities:

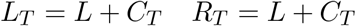

where 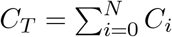

Using values of *L*_*T*_ and *R*_*T*_ in equation (43) we get

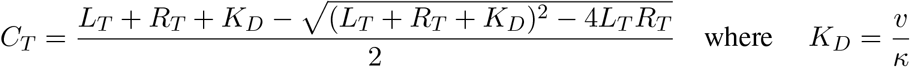

Let

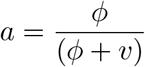

Hence, at equilibrium

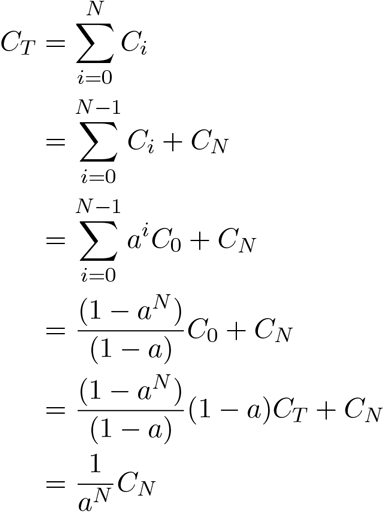

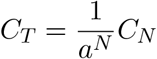

Hence

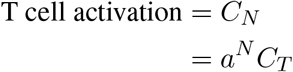

#### Calculation for *E*_max_

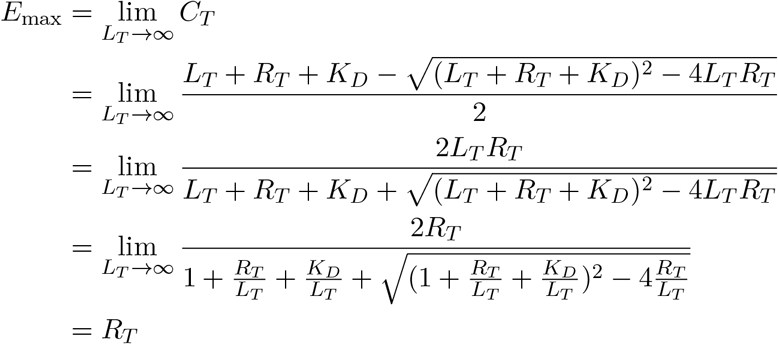

So *E*_max_ = *a*^*N*^*R*_*T*_

#### Calculation for *EC*_50_

At half the maximal response, T cell activation =

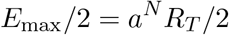

which implies

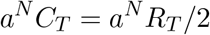

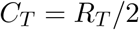

Since, *L*_*T*_ = *L* + *C*_*T*_ *R*_*T*_ = *L* + *C*_*T*_ Using this equation in equation (43)

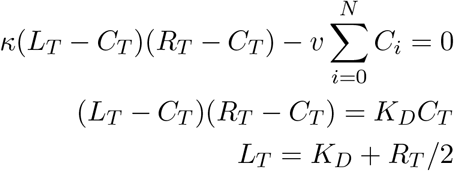

### C.3 Kinetic proofreading with limited signaling

It is an extension of the kinetic proofreading model that proposes that when a TCR has reached signalling competent state *C*_*N*_, the bound TCR transits to a non signalling state *C*_*N*+1_ with rate *ξ*.

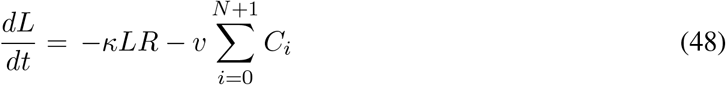

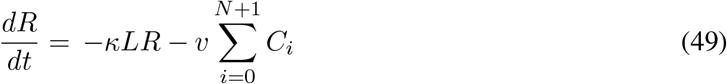

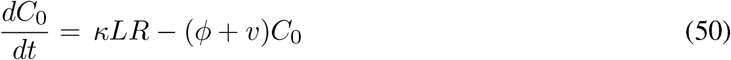

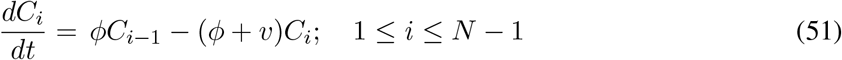

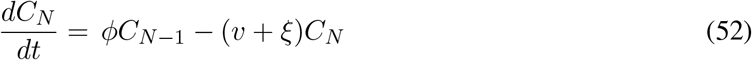

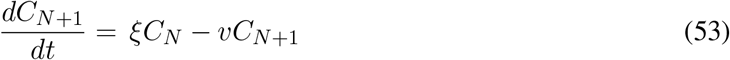

Here 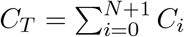, and T cell activation is given by *C*_*N*_.

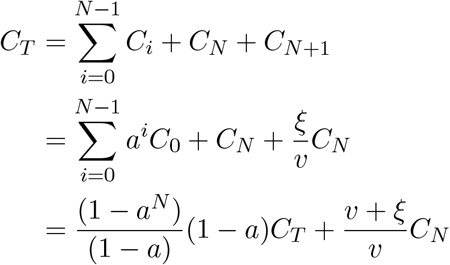

Hence

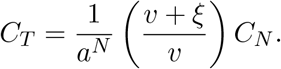

and

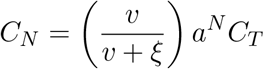

#### Calculating the *E*_max_

Similar to the case of kinetic proofreading model, it can be shown that when *L*_*T*_ → ∞ *C*_*T*_ → *R*_*T*_ Hence

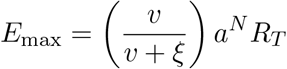

#### Calculating the *EC*_50_

At half the maximal response, T cell activation

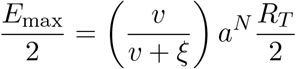

which implies

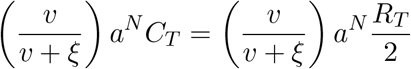

Hence

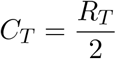

Since, *L*_*T*_ = *L* + *C*_*T*_ *R*_*T*_ = *R* + *C*_*T*_ Using this equation in (48)

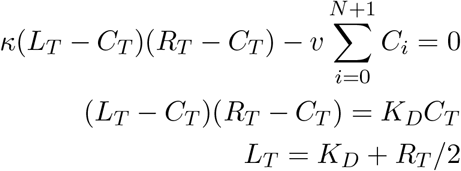

### C.4 Kinetic proofreading with sustained signaling

It is another modification of the kinetic proofreading model, according to this model, T-cell receptors (TCRs) in the signaling-competent state *C*_*N*_, persist in signaling for a certain duration (T) after the unbinding of pMHC. Subsequently, they return to the basal state (T) with a rate of *λ*.

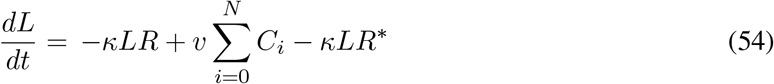

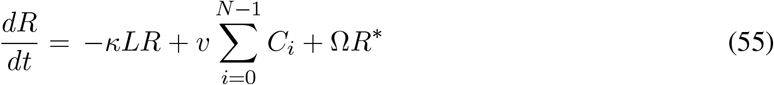

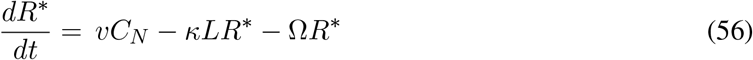

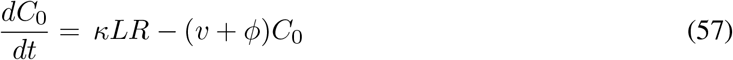

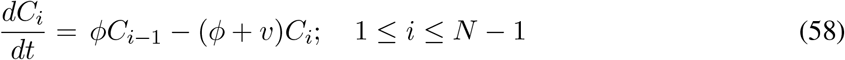

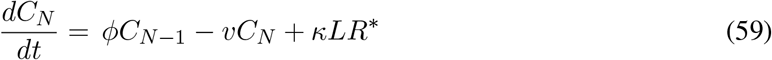

The conservation equations are:

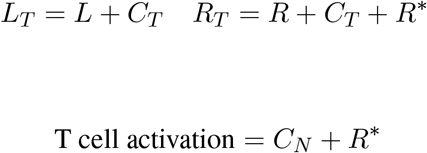

From equation (56) at the steady state

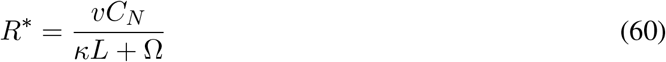

Also substituting (60) into (55) at steady state we get

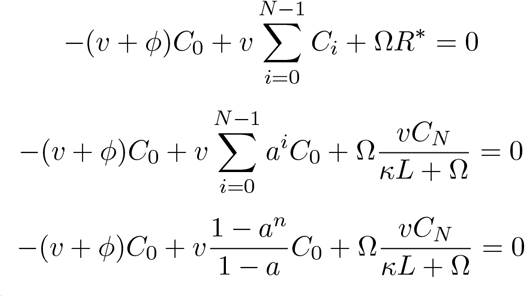

upon rearrangement and solving we get

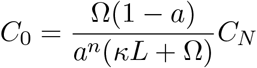

Also,

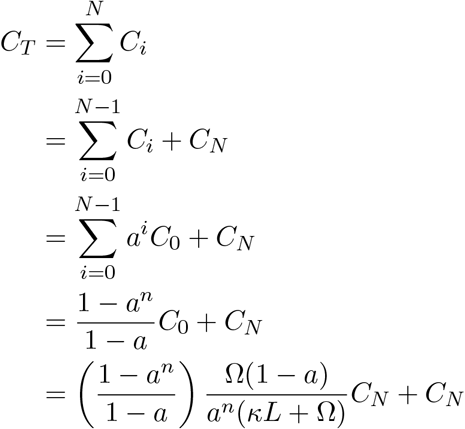

Solving this we get:

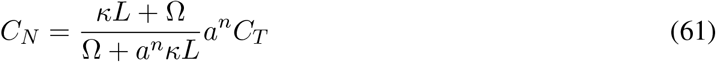

Hence from equation (60) and (61), T cell activation is given by

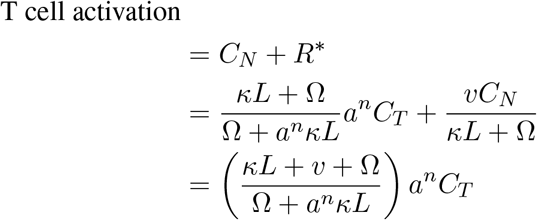

#### Calculating the *E*_max_

Similar to the case of kinetic proofreading model, it can be shown that when *L*_*T*_ → ∞ *C*_*T*_ → *R*_*T*_ where *C*_*T*_ is given by:

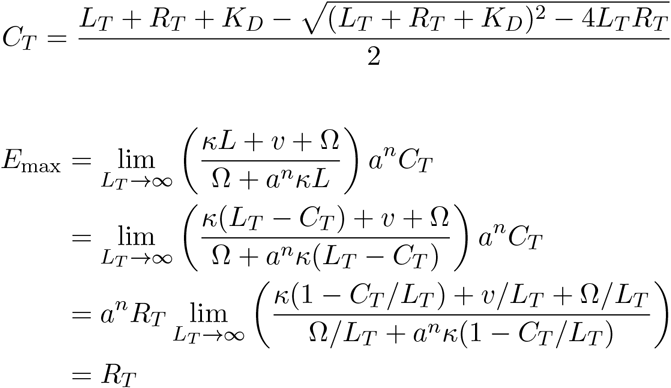

Hence *E*_max_ = *R*_*T*_

#### Calculating the *EC*_50_

At half the maximal response

T cell activation

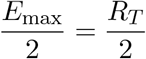

which implies

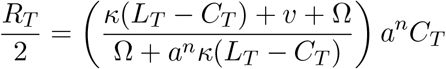

Rearranging and solving for *L*_*T*_ we get:

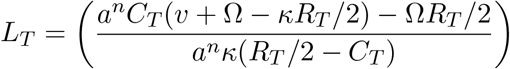

### C.5 Kinetic proofreading with negative feedback

Kinetic Proofreading with Negative Feedback extends the KPR scheme by introducing the notion that the rates of complex formation in the activation chain can be modulated at intermediate stages and/or within the final signaling state *C*_*N*_. This modulation is achieved through a single negative feedback mechanism mediated by Src homology 2 domain phosphatase-1 (SHP-1).

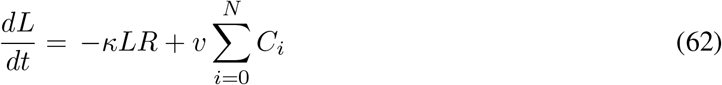

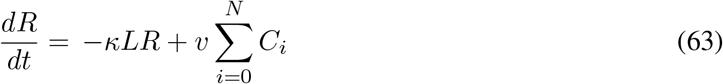

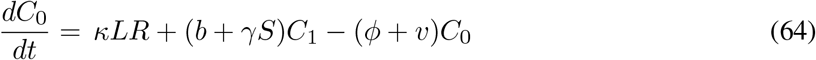

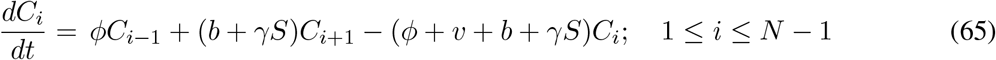

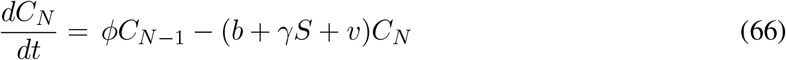

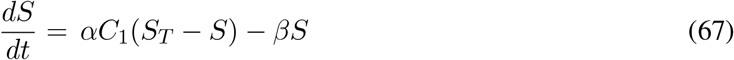

The conservation equations are:

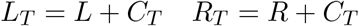

Here 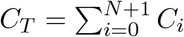, and T cell activation is given by *C*_*N*_.

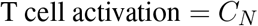

Now *C*_*N*_ can be expressed in form of *C*_*T*_ as in eq (4.17) in [25],

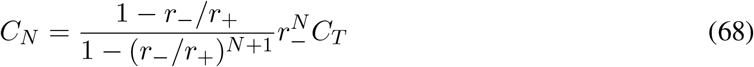

where

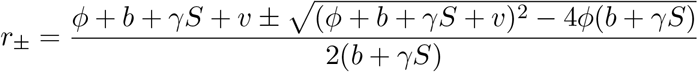

Therefore, the cell activation is given by

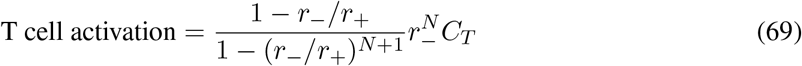

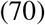

#### Calculation for *E*_max_

Similar to the case of kinetic proofreading model, it can be shown that when *L*_*T*_ → ∞ *C*_*T*_ → *R*_*T*_ where *C*_*T*_ is given by:

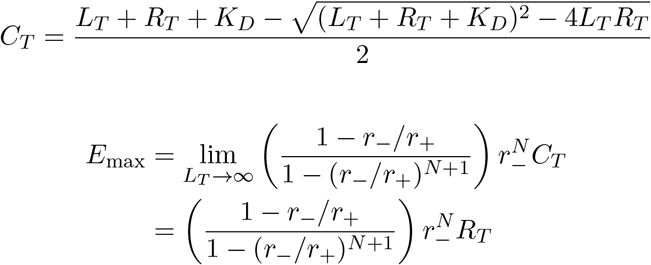

#### Calculation for *EC*_50_

At half the maximal response

T cell activation

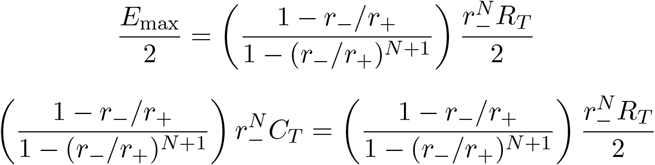

which implies

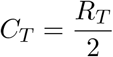

Since, *L*_*T*_ = *L* + *C*_*T*_ *R*_*T*_ = *L* + *C*_*T*_, using this equation in equation 62

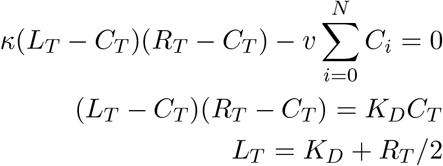

### C.6 Kinetic proofreading with stabilizing activation chain

In this model a pMHC ligand (L) binds to a TCR receptor (R) to form a pMHC-TCR complex *C*_0_, which undergoes series of chemical modifications to reach signalling competent state *C*_*N*_.

This stabilization/destabilization of the *C*_*i*_ complexes and variation in the time taken for activation, are articulated by variations in the values of corresponding rate constants *v*(*i*); (*i* = 0, 1, …, *N*) and *ϕ*(*i*); (*i* = 0, 1, …, *N* − 1) as the proofreading progresses.

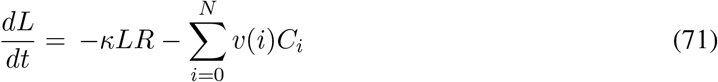

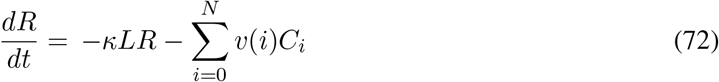

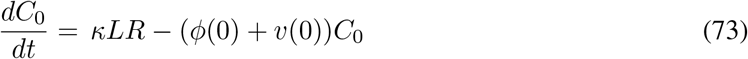

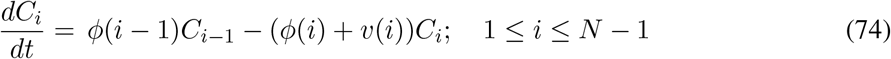

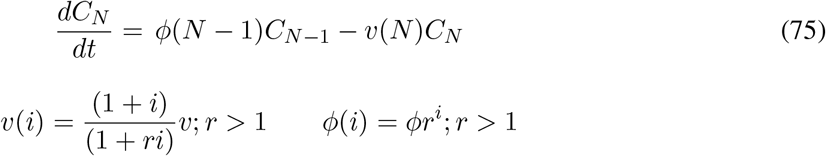

where *v*(0) = 1.5 and *ϕ*(0) = 1.3.

In this model for simplicity *v*(0) is denoted as *v*(= 1*/τ*) which is dissociation time, and *ϕ*(0) denotes the propagation rate for first step *C*_0_ → *C*_1_. Now for next steps these rates are given by

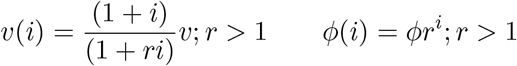

The conservation equations are:

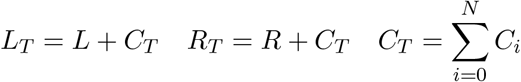

Here we have from [30]

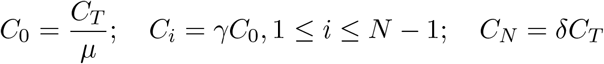

where

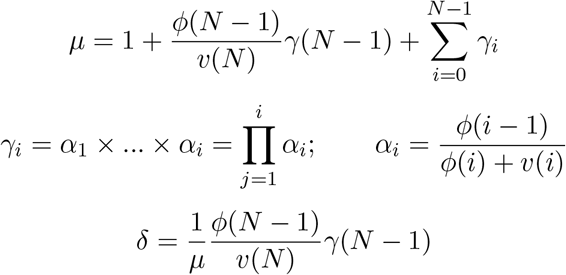

and where *C*_*T*_ is the number of bound receptors or ligands 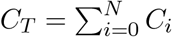 which is given by

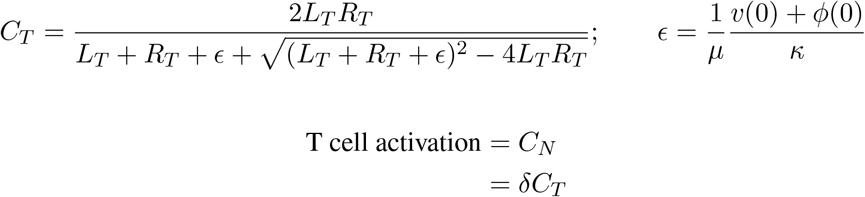

#### Calculating the *E*_max_

Similar to the case of kinetic proofreading model, it can be shown that when *L*_*T*_ → ∞ *C*_*T*_ → *R*_*T*_ Hence

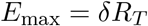

#### Calculating the *EC*_50_

At half the maximal response

T cell activation

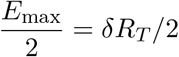

which implies

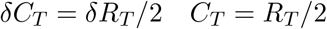

Since

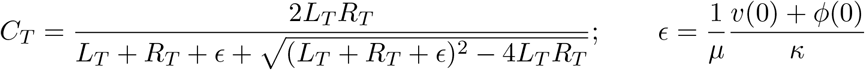

Substituting *C*_*T*_ = *R*_*T*_ */*2 in above equation and solving for *L*_*T*_ we get

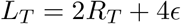

Hence potency

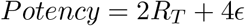

### C.7 Kinetic proofreading with incoherent feed forward loop

Although the actual model [31] considered the KPR with limited signalling combined with incoherent feed forward loop. We first considered the KPR with incoherent feed forward loop only for the plotting.

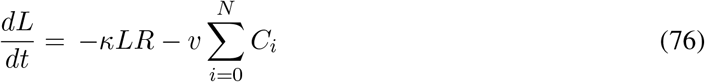

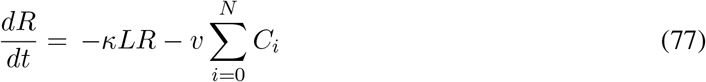

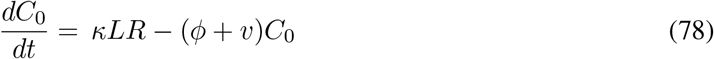

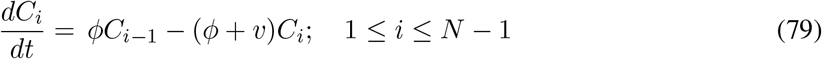

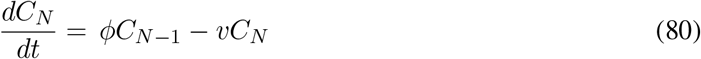

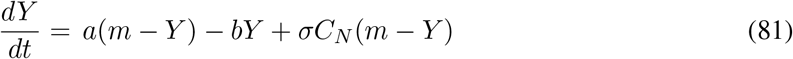

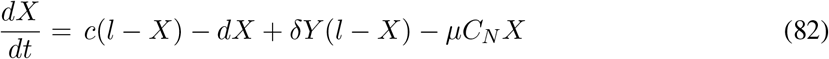

where 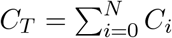, and T cell activation is given by X.

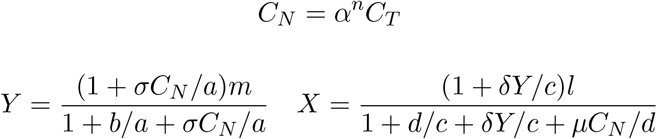

#### Calculating the *E*_max_

Similar to the case of kinetic proofreading model, it can be shown that when *L*_*T*_ → ∞ *C*_*T*_ → *R*_*T*_

Hence

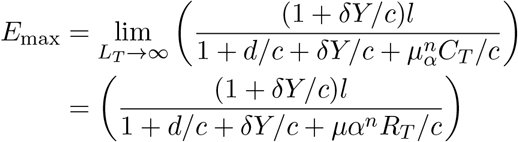

#### Calculating the *EC*_50_

At half the maximal response

T cell activation

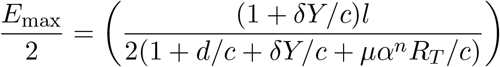

which implies

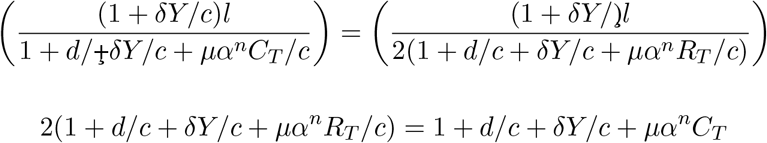

Rearranging and solving for *C*_*T*_

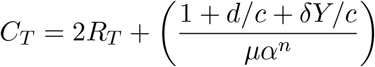

Since, *L*_*T*_ = *L* + *C*_*T*_ *R*_*T*_ = *R* + *C*_*T*_ Using this equation in (76)

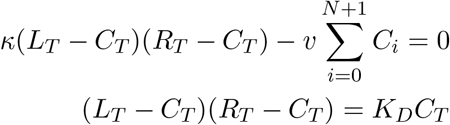

Using value of *C*_*T*_ in above equation and solving. we get:

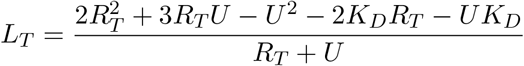

where

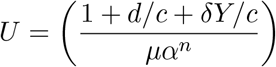

### C.8 Kinetic proofreading with limited signaling and incoherent feed forward loop

This model is an extension of KPR with incoherent feed forward loop, it has been assumed that signalling is limited and after after reaching the signalling competent state the bound TCR transits to non-signalling transit state.

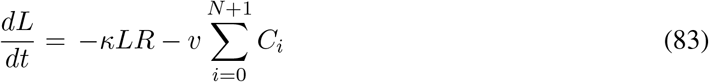

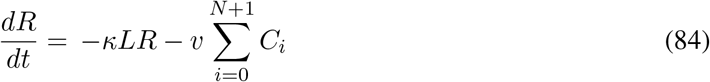

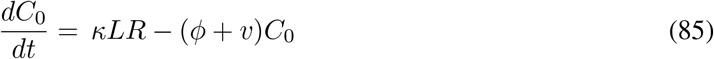

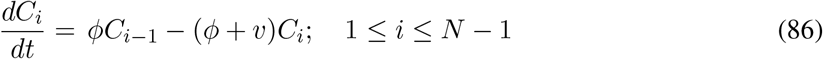

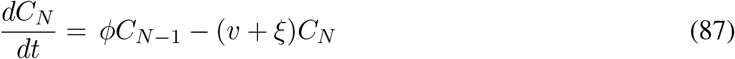

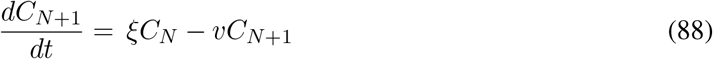

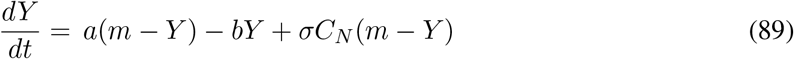

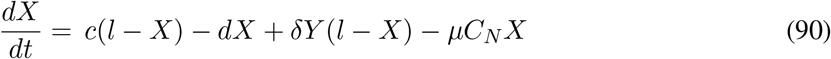

where 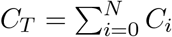, and T cell activation is given by X.

Here

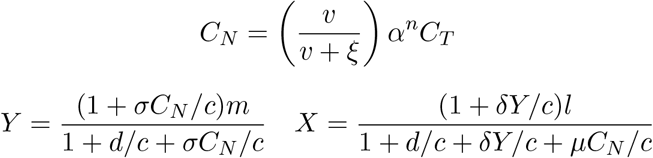

#### Calculating the *E*_max_

Similar to the case of kinetic proofreading model, it can be shown that when *L*_*T*_ → ∞ *C*_*T*_ → *R*_*T*_

Hence

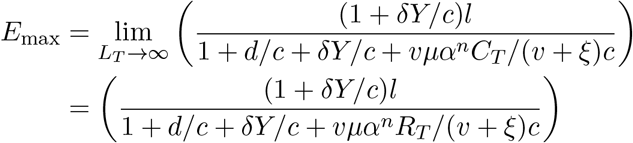

#### Calculating the *EC*_50_

At half the maximal response

T cell activation

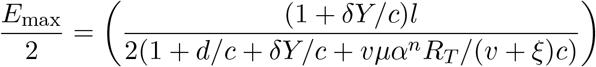

which implies

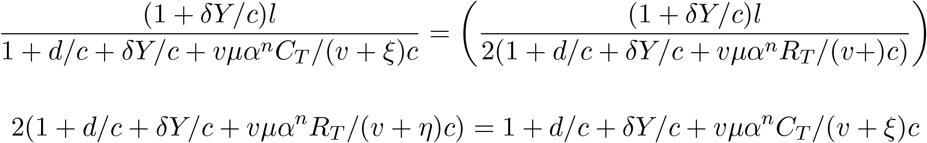

Rearranging and solving for *C*_*T*_

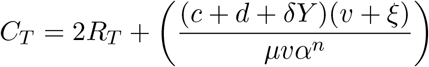

Since, *L*_*T*_ = *L* + *C*_*T*_ *R*_*T*_ = *R* + *C*_*T*_, using this equation in (83)

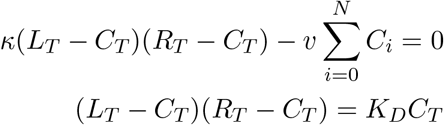

Using value of *C*_*T*_ in above equation and solving. we get:

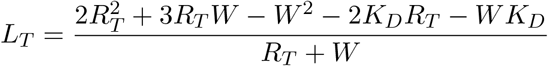

where

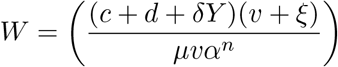

### C.9 Kinetic proofreading with Limited and Sustained Signaling

It as a combination of two models KPR with limited signaling and KPR with sustained signaling.

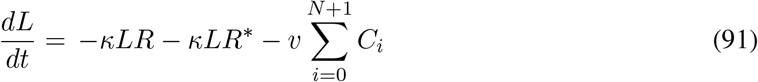

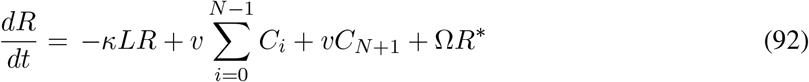

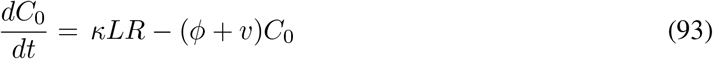

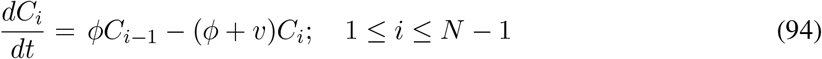

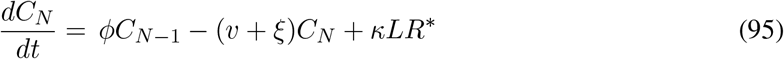

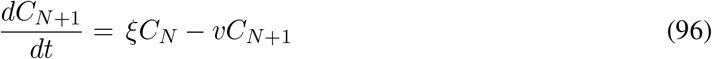

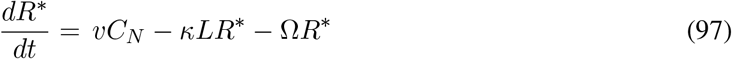

The conservation equations are:

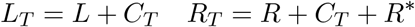

where 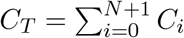

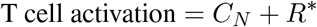

From (97) at the steady state

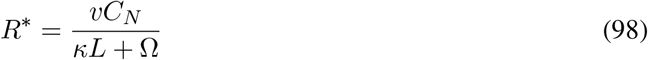

Now, using it and (93) at a steady state; for (92) at a steady state we have

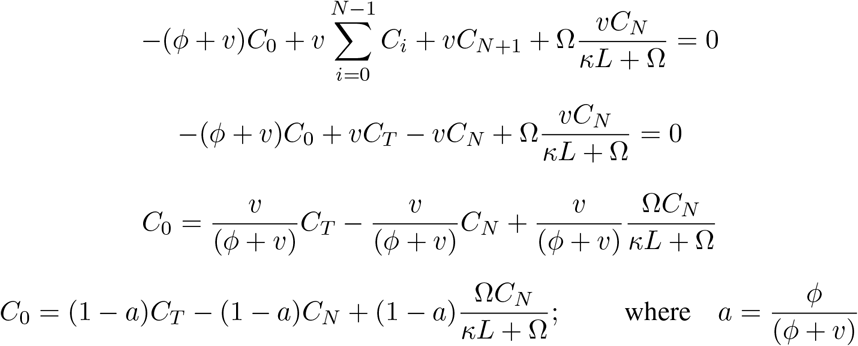

As *C*_*T*_ is given by:

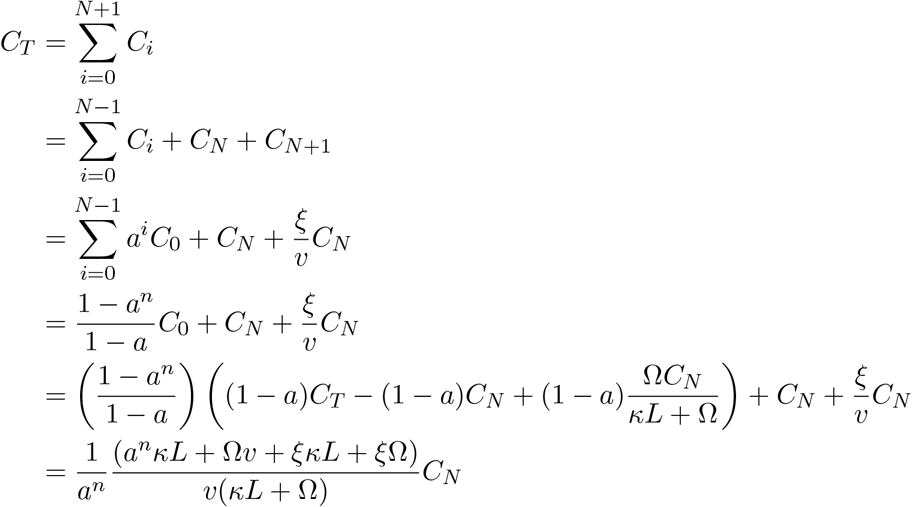

Which implies *C*_*N*_ is given by:

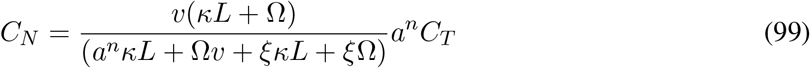

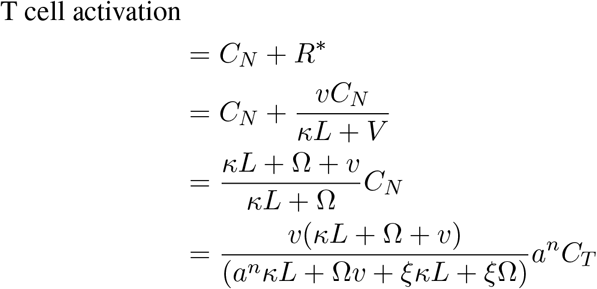

#### Calculating the *E*_max_

Similar to the case of kinetic proofreading model, it can be shown that when *L*_*T*_ → ∞ *C*_*T*_ → *R*_*T*_ where *C*_*T*_ is given by:

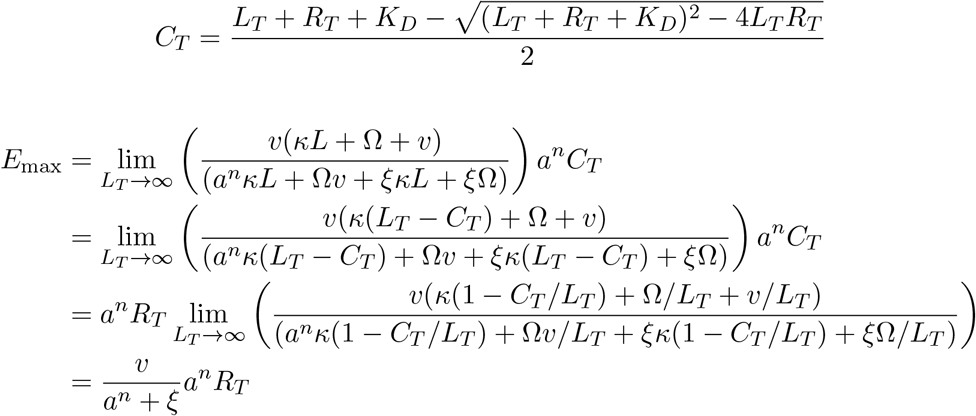

#### Calculating the *EC*_50_

At half the maximal response

T cell activation

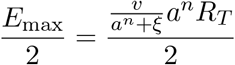

which implies

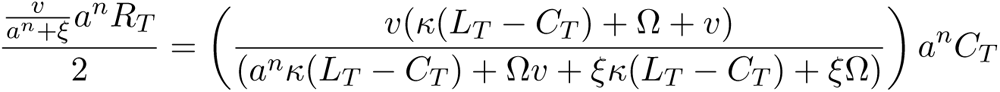

Rearranging and solving for *L*_*T*_ we get:

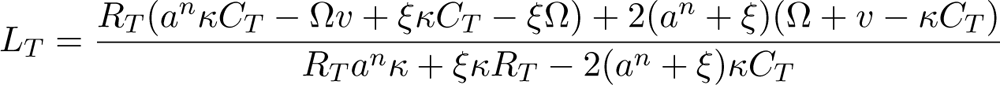

